# Broad transcriptomic dysregulation across the cerebral cortex in ASD

**DOI:** 10.1101/2020.12.17.423129

**Authors:** Jillian R. Haney, Brie Wamsley, George T. Chen, Sepideh Parhami, Prashant S. Emani, Nathan Chang, Gil D. Hoftman, Diego de Alba, Gaurav Kale, Gokul Ramaswami, Christopher L. Hartl, Ting Jin, Daifeng Wang, Jing Ou, Ye Emily Wu, Neelroop N. Parikshak, Vivek Swarup, T. Grant Belgard, Mark Gerstein, Bogdan Pasaniuc, Michael J. Gandal, Daniel H. Geschwind

## Abstract

Classically, psychiatric disorders have been considered to lack defining pathology, but recent work has demonstrated consistent disruption at the molecular level, characterized by transcriptomic and epigenetic alterations.^1–3^ In ASD, upregulation of microglial, astrocyte, and immune signaling genes, downregulation of specific synaptic genes, and attenuation of regional gene expression differences are observed.^1,2,4–6^ However, whether these changes are limited to the cortical association areas profiled is unknown. Here, we perform RNA-sequencing (RNA-seq) on 725 brain samples spanning 11 distinct cortical areas in 112 ASD cases and neurotypical controls. We identify substantially more genes and isoforms that differentiate ASD from controls than previously observed. These alterations are pervasive and cortex-wide, but vary in magnitude across regions, roughly showing an anterior to posterior gradient, with the strongest signal in visual cortex, followed by parietal cortex and the temporal lobe. We find a notable enrichment of ASD genetic risk variants among cortex-wide downregulated synaptic plasticity genes and upregulated protein folding gene isoforms. Finally, using snRNA-seq, we determine that regional variation in the magnitude of transcriptomic dysregulation reflects changes in cellular proportion and cell-type-specific gene expression, particularly impacting L3/4 excitatory neurons. These results highlight widespread, genetically-driven neuronal dysfunction as a major component of ASD pathology in the cerebral cortex, extending beyond association cortices to involve primary sensory regions.

## Main Text

### Transcriptomic changes across the cerebral cortex in ASD

Similar to other neuropsychiatric disorders, the risk for autism spectrum disorder (ASD) involves substantial genetic liability, which is profoundly complex and heterogeneous.^7,8^ Despite this causal heterogeneity, molecular profiling studies consistently show common patterns of shared transcriptomic and epigenetic dysregulation in the majority of ASD cases in frontal and temporal association cortex.^1–3,5^ Whether this represents focal, regional, or more generalized dysfunction is not known. To address this question cortex-wide, we conducted strand-specific RNA-sequencing (RNA-seq) to identify gene and isoform (transcriptomic) changes in 725 samples across 11 brain regions spanning all four cortical lobules (frontal, parietal, temporal, and occipital), from 49 subjects with idiopathic ASD and 54 matched neurotypical controls (**Fig. 1a**, **Methods**, **Supplementary Table 1**, and **Extended Data Fig. 1-3**). Previous work using gene expression microarrays and RNA-seq identified gene co-expression modules representing specific pathways differentially expressed in ASD frontal and temporal cortices.^4,5^ The number of samples profiled here is more than five times greater than these prior studies, so we first used this multi-region RNA-seq resource to replicate and extend these previous findings. We observed widespread dysregulation across all 11 cortical regions that replicated the previously identified patterns of dysregulation in temporal and frontal cortex (**Fig. 1b**, **Methods**, **Supplementary Table 2**, **Extended Data Fig. 3**). However, the magnitude of effect varied across regions, with the primary visual cortex (V1; Brodmann Area (BA) 17) exhibiting the greatest degree of dysregulation, followed by parietal cortex (BA7) and posterior superior temporal gyrus (BA 41/42/22) both in terms of fold changes and the number of genes differentially expressed (**Fig. 1b**, **Extended Data Fig. 3**). To show that this was not due to regional variation in sample sizes, we performed permutation testing, which indicated that this increased signal was not biased by regional sample size differences (**Methods**, **Supplementary Table 2**).

**Figure 1.**
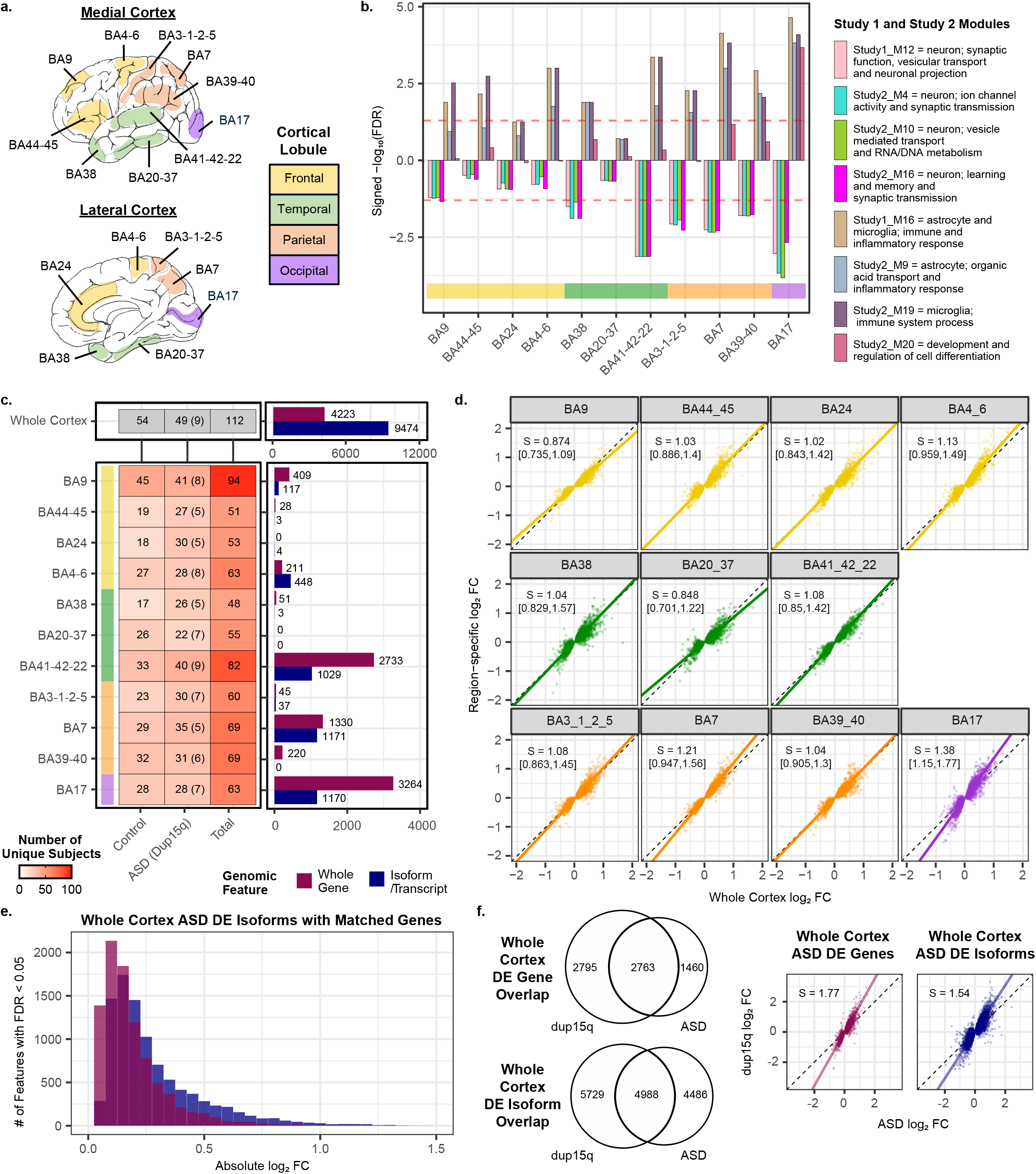
ASD Transcriptomic Differences Across 11 Cortical Regions. **a.** Human cortical Brodmann Areas (BA) with cortical lobules indicated. Cortical lobule colors are consistent throughout the figure. **b.** Dysregulation of previously identified co-expressed gene modules (Study 1: Voineagu et al., *Nature* 2011; Study 2: Parikshak et al., *Nature* 2016)4-5 across cortical regions. Red dashed line marks FDR < 0.05. **c.** Unique subjects by region and diagnosis (left), with number of differentially expressed (DE; linear mixed model FDR < 0.05) genes and isoforms (right) across the whole cortex (top) or within individual regions (bottom). Gene and isoform colors are consistent throughout this figure. **d.** log2 fold change (FC) of individual regions compared to the whole cortex log2 FC for the 4,223 whole cortex DE genes. Slope (S, with 95% confidence interval in brackets) is calculated with principal components regression. **e.** For the whole cortex DE isoforms, a histogram of all isoforms DE across the whole cortex in ASD with their matched genes. **f.** Left: venn diagrams depicting the number of genes and isoforms DE across the whole cortex in dup15q samples compared to idoipathic ASD samples. Right: for the ASD whole cortex DE genes (left) and isoforms (right), the idiopathic ASD whole cortex log2 FC compared to the dup15q whole cortex log2 FC. Slope (S) is calculated with principal components regression.

Given that qualitatively similar transcriptomic changes were observed across regions (**Fig. 1b**), we next combined all regions to increase our statistical power to detect previously unrecognized differentially expressed (DE) genes and isoforms. We used a linear mixed model framework to control for individual effects and identify changes across all 11 regions examined as well as within individual regions, separately (**Fig. 1c**, **Methods**, **Supplementary Table 3**, **Extended Data Fig 4**). We found 4,223 genes and 9,474 isoforms (FDR < 0.05) DE across all cortical regions, a notable increase compared to previous analyses (**Fig. 1c**, **Extended Data Fig. 3**). We again observed the greatest signal in BA17, and 59% of DE genes in BA17 alone overlapped with what was observed globally (**Supplementary Table 2**, **Extended Data Fig. 4**). Additionally, DE gene effect sizes in BA17 and BA7 were the highest in magnitude, more than other regions assessed (**Fig. 1d**, **Methods**). In comparing DE genes and isoforms across all regions, we found both conserved and distinct dysregulation (**Extended Data Fig. 4**, **Supplementary Table 2**). Notably, as previously observed in frontal and temporal cortex^1^ we observed that DE isoforms exhibited greater effect size changes in ASD than their matched genes (**Fig. 1e**, **Supplementary Table 2**, **Extended Data Fig. 4**).

We next evaluated differential gene and isoform expression in an additional 83 pan-cortical samples from 9 subjects with dup15q syndrome, a rare genetic disorder with high penetrance for ASD, which previously was shown to strongly parallel changes in idiopathic ASD in frontal and temporal cortex, but with greater magnitude of effect.^5^ We replicated these previous results broadly across the cortical regions examined, finding substantial overlap in transcriptomic changes between dup15q and idiopathic ASD and with dup15q exhibiting a greater magnitude of dysregulation overall (**Fig. 1f**, **Supplementary Table 2**, **Extended Data Fig. 4**). BA17 also exhibited the greatest number of DE genes in dup15q (**Extended Data Fig. 4**). These results demonstrate that the molecular pathology shared by this genetic form of ASD and idiopathic ASD is widespread across distinct regions of the cortex, and that some commonalities in regional variance of effect exist, both impacting sensory in addition to association cortex.

### Broad attenuation of transcriptomic regional identity

We previously observed an attenuation of typical gene expression differences between two regions, frontal and temporal lobe in ASD,^4,5^ which we refer to here as an “Attenuation of Transcriptomic Regional Identity” (ARI). To assess whether this was a broader phenomenon, we systematically contrasted all unique pairs of 11 cortical regions (55 comparisons in all) using a conservative statistical approach to account for differences in sample size across regions, while correcting stringently for multiple comparisons (**Fig. 2a**, **Methods**). We further validated the identified transcriptomic regional identity patterns in our control samples with those from an external data source, the Allen Brain Atlas^9^ (**Supplementary Table 4**, **Extended Data Fig. 5**, **Methods**). Ten pairs of regions exhibited significantly greater ARI patterns in ASD compared to controls, with an additional 31 out of the 55 pairs of regions exhibiting a trend towards attenuation in ASD (**Fig. 2b**, **Supplementary Table 4**, **Extended Data Fig. 5**, **Methods**). These results provide evidence in support of widespread ARI across the cerebral cortex in ASD for the first time, across both gene and isoform levels (**Extended Data Fig. 5**). Additionally, we observed a regional anterior - posterior gradient, with nine of the ten region pairs exhibiting significant ARI in ASD containing either BA17 or BA39-40 (**Fig. 2c-d**). Notably, BA17 was also one of the regions with the largest case-control differences in gene expression. To determine how gene expression changes were dispersed across regions in these pairs, we used a conservative filtering process to identify individual genes exhibiting ARI (**Methods**, **Supplementary Table 4**). Although these genes were widely dysregulated, the posterior regions BA17 and BA39-40 exhibited the greatest changes (**Fig. 2c-d**, **Extended Data Fig. 6**). ARI genes were also comparably disrupted in the dup15q samples (**Extended Data Fig. 6**), suggesting that transcriptomic regional identity attenuation in the cerebral cortex is shared across heterogenous forms of ASD.

**Figure 2.**
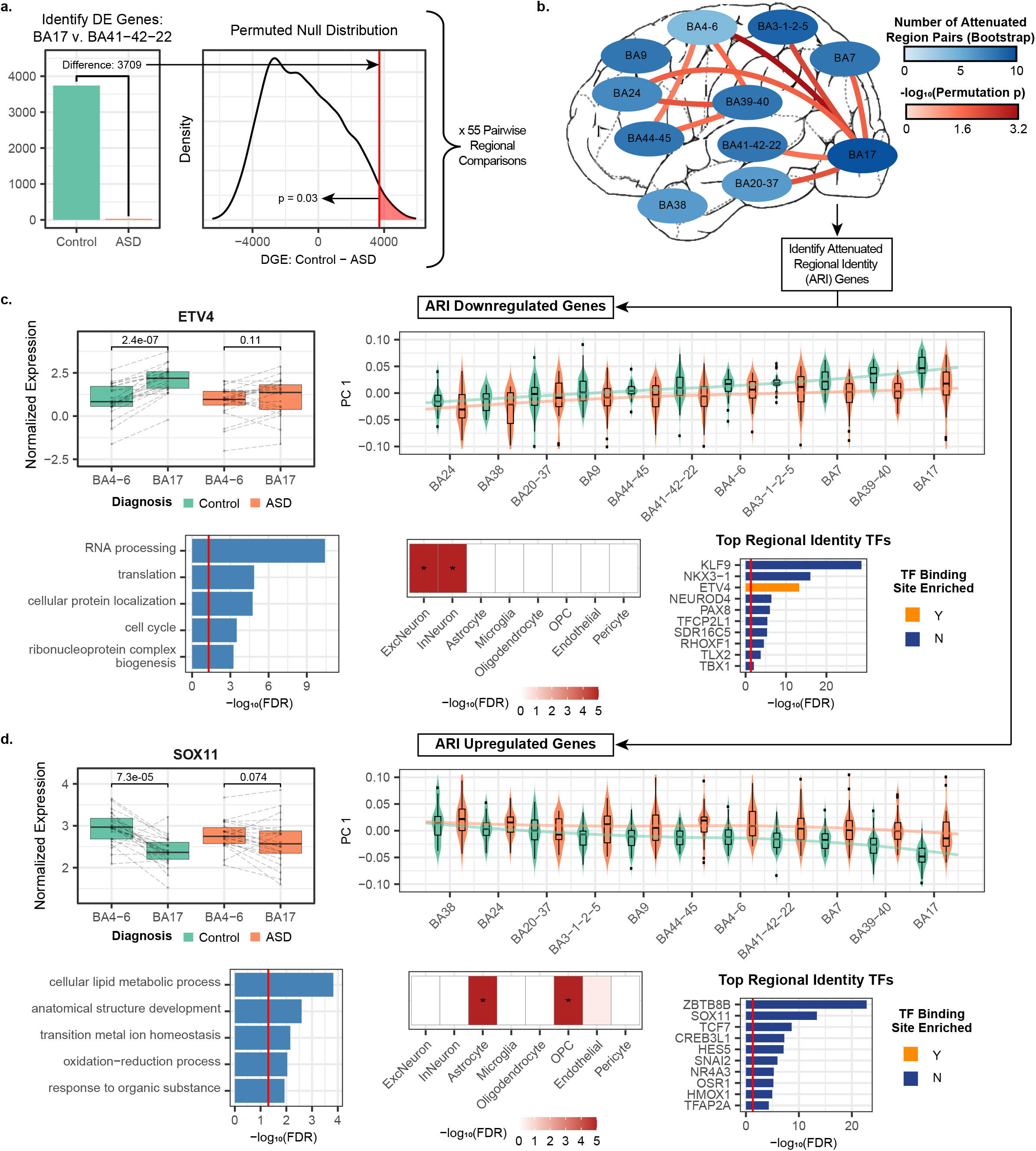
Transcriptomic Regional Identity Attenuation in ASD. **a.** Methods overview for identifying statistically significant differences in transcriptomic regional dentity in ASD. The regional comparison of BA17 v. BA41-42-22 is used here as an example. Number of DE genes between regions is calculated in controls and ASD samples (left). A permuted null distribution is then used to determine the significance of the difference in DE genes between controls and ASD samples (right). **b.** Regional comparisons with attenuation of transcriptomic regional identity in ASD with permutation p < 0.05 are connected with a bar (red scale), and a general trend towards cortex-wide attenuation is summarized by region color (blue scale; 0=no pairs exhibiting attenuation in ASD, 10=every pair exhibits attenuation in ASD). For region color, a regional pair is considered attenuated if it contains less DE in ASD compared to controls as measured with a bootstrap approach (**Methods**). Attenuated regional identity (ARI) genes are extracted from regional comparisons with permutation p < 0.05 (**Methods**). **c-d.** Overview of ARI downregulated (**c.**) and upregulated (**d.**) genes. Top left, select attenuated transcription factors in BA17 and BA4-6. Lines link paired samples from the same subject, and the paired Wilcoxon signed-rank test p-value is plotted above boxplots. Top right, PC 1 of ARI genes across all regions. Bottom left and bottom center, gene ontology and cell-type enrichment, respectively. Bottom right, top 10 attenuated transcription factors (TFs), where FDR is representative of how well these TFs distinguish BA17 and BA39-40 from the other nine cortical regions assessed here in controls (Methods). Enrichment for transcription factor binding sites is also depicted (Bonferroni-corrected p-value < 0.05 required for enrichment). Please see the Supplementary Methods for a description of all boxplots included in figures

To identify the biological processes contributing to ARI gene dysregulation in ASD, we grouped together all of the ARI genes that were either downregulated (1,881 genes) or upregulated (1,695 genes) with a pronounced posterior effect in ASD (**Methods**). The downregulated set of ARI genes showed broad enrichment for neuronal cell-type-specific markers and RNA processing pathways, and contained many transcription factors (**Fig. 2c**, **Supplementary Table 4**). The upregulated ARI genes also contained many transcription factors and were enriched for oligodendrocyte progenitor cell (OPC) and astrocyte cell-type markers along with metabolic and development pathways. ARI gene dysregulation was further characterized by subsequent co-expression network analysis, which further refined the topology and pathways involved.

### Refining disrupted gene co-expression networks in ASD

We next used weighted gene correlation network analysis (WGCNA)^10^ across all samples to partition genes into co-expression modules capturing potentially shared biological functions or regulation (**Methods**). We identified a total of 35 gene modules, of which 9 were downregulated and 15 were upregulated in ASD (**Supplementary Table 5-6**, **Extended Data Fig. 7**). We further generated networks using isoform-level quantifications, identifying 61 isoform modules. Of these, 39 were distinct from the gene modules, with 5 downregulated and 9 upregulated in ASD (**Supplementary Table 5-6**, **Extended Data Fig. 8**). In total, 38 gene and isoform modules were dysregulated in at least one region in ASD. Most of these fell into two broad groups - either dysregulated (1) cortex-wide with comparable magnitude across regions, or (2) with significantly variable magnitude across regions. Again, dup15q effects were similar to ASD effects, but were greater in magnitude (**Extended Data Fig. 7-8**, **Supplementary Table 6**).

#### Cortex-wide dysregulation observed for ASD risk genes

Eighteen gene and isoform modules exhibited a consistent pattern of dysregulation in ASD across all cortical regions assessed (linear mixed model, FDR < 0.05; **Fig. 3a**, **Extended Data Fig. 7-8**, **Supplementary Table 6**). These include GeneM9, an upregulated neuronal module with a significant enrichment for non-coding genes; GeneM32, a strongly upregulated reactive astrocyte module with the greatest overall magnitude of dysregulation; and GeneM24, a downregulated module enriched for endothelial and pericyte marker genes which are involved in blood-brain-barrier functions (**Fig. 3b, Extended Data Fig. 7, Supplementary Table 6**). These modules replicate previous findings of neuronal upregulation, astrocyte reactivity, and BBB disruption in ASD,^1,4–6^ but extend these findings by demonstrating that these processes are widespread across the cerebral cortex.

**Figure 3.**
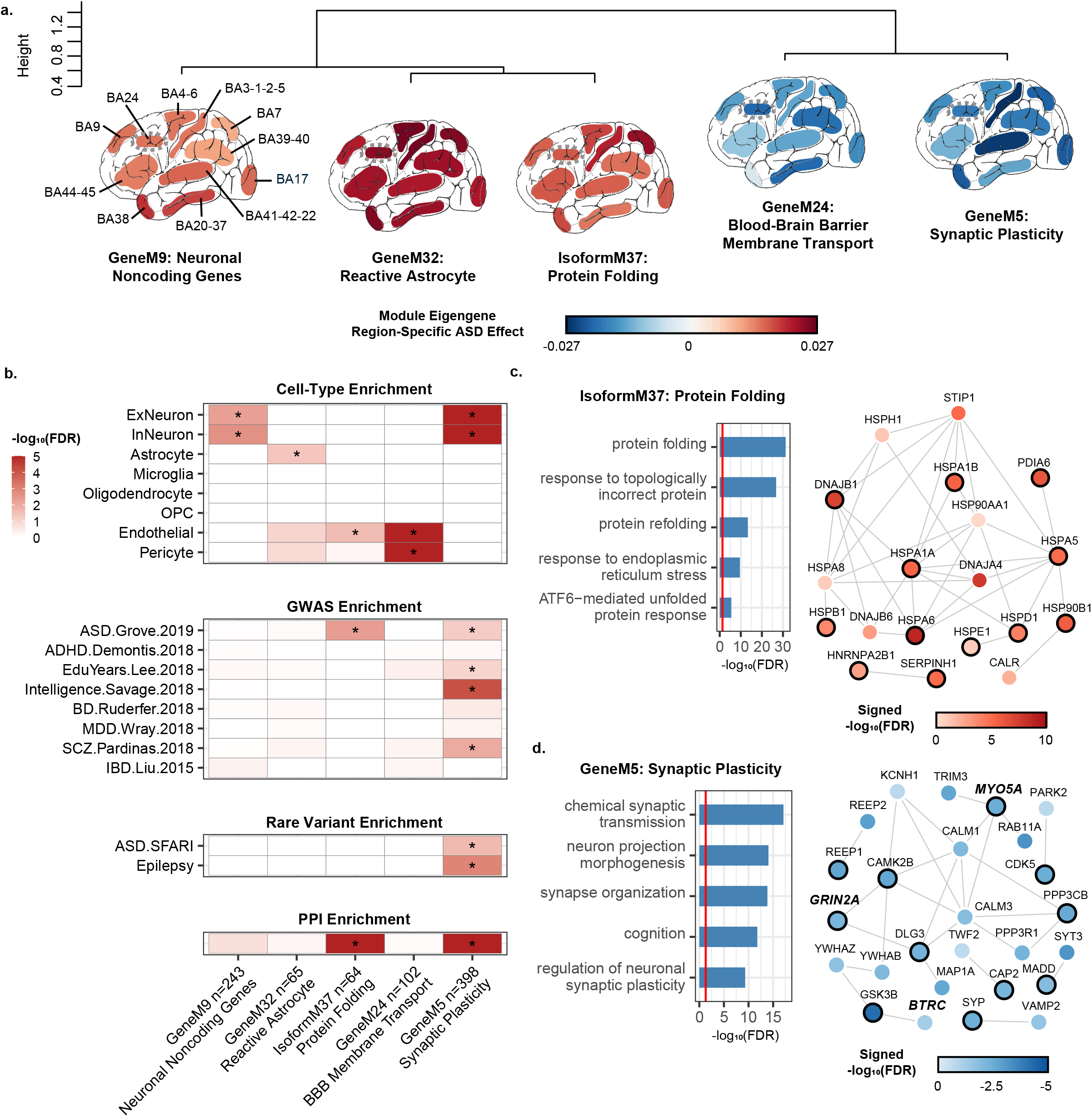
Co-expression Network Analysis Characterizes Cortex-wide Dysregulation of ASD Risk Genes. **a.** Average linkage hierarchical clustering of the biweight midcorrelation of the top 5 most dysregulated gene and isoform co-expression module eigengenes (first principal component of the module) with regionally-consistent patterns of ASD dysregulation. The module eigengene ASD effect is indicated for each cortical region examined (**Methods**). **b.** −log10(FDR) for cell-type, GWAS, rare variant, and protein-protein interaction (PPI) enrichment for the modules depicted in **a**. ‘*’ indicates a significant enrichment (FDR < 0.05 for cell-type, rare variant, and PPI enrichment, and FDR < 0.1 for GWAS enrichment). ‘n’ indicates the number of genes/isoforms in each module. **c-d.** For ASD GWAS enriched modules Isoform M37 (**c**) and GeneM5 (**d**), top gene ontology terms (left) and hub genes (module genes within the top 20 genes with the highest correlation with the module’s eigengene) that participate in a protein-protein interaction (PPI) with any other module gene are depicted along with their PPI partners (right). Node color is the signed −log10(FDR) of the whole cortex ASD effect, edges denote direct PPIs, and hub genes are indicated with a black outline. SFARI database14 gene names are in bold italic.

Two modules - GeneM5 and IsoM37 - demonstrated cortex-wide dysregulation along with significant enrichment for ASD-associated common genetic variation (**Fig. 3b-d**).^11^ GeneM5 is down-regulated in ASD, contains many neuronal genes involved in synaptic plasticity, and significantly overlaps with the synaptic module CTX.M16 previously identified by Parikshak et al.^5^ (**Fig. 3a**, **Fig. 3d**, **Supplementary Table 5-6**). In addiiton to common genetic variation, GeneM5 is also significantly enriched for genes containing rare *de novo* protein disrupting mutations associated with ASD, including the high-confidence risk genes *GRIN2A*, *MYO5A*, and *BTRC*^12^ (**Supplementary Table 5-6**, **Methods**). This demonstrates another point of convergence of rare and common risk variants on shared biological processes in ASD.^39^ GeneM5 is enriched in cortical lower layer 4-6 excitatory neuron cell-type markers (**Extended Data Fig. 7**),^13^ identifying them as a point of convergence for rare and common genetic risk in ASD. Finally, IsoM37 is enriched for ASD common genetic risk variants (but not rare mutations), is upregulated in ASD, and contains genes involved in protein folding (**Fig. 3a**, **Fig. 3c**, **Supplementary Table 6**). To our knowledge, this is the first report of an upregulated ASD transcriptomic signature that is associated with known ASD risk variants.

#### Magnitude of effect parallels anterior-posterior gradients

In addition to observing profound cortex-wide dysregulation in ASD, we found 13 modules that exhibited their most pronounced ASD effect in BA17, as measured against a permuted distribution containing all regions (**Extended Data Fig. 7**, **Supplementary Table 6**, **Methods**). Of these, 12 showed significant enrichment for ARI genes (half up-regulated and half down-regulated in ASD) and all 13 had anterior - posterior gradients of expression in neurotypical samples, indicating that these modules contribute to transcriptomic regional identities that are observed in neurotypical controls, but attenuated in ASD (**Extended Data Fig. 7**, **Supplementary Table 6**). Six of these modules were more highly expressed in posterior regions in neurotypical subjects and were observed to be downregulated in ASD across the cortex (**Fig. 4a**, **Extended Data Fig. 7**, **Supplementary Table 6**). These include GeneM23, an oligodendrocyte-specific module consisting of genes important for organelle regulation and intracellular restructuring; GeneM14, a neuronal module that contains genes involved in neurite morphogenesis and is also strongly downregulated in BA41-42-22; and GeneM3, a neuronal module enriched for energy generation and neuronal processes that are highly energy dependent, such as vesicle transport (**Fig. 4b-c**, **Supplementary Table 5-6**). GeneM3 is also significantly enriched for cell-type markers specific to layer 4-5 excitatory neurons (**Extended Data Fig. 7**).^13^ The next four modules were more highly expressed in anterior regions in neurotypical subjects and exhibited cortex-wide upregulation in ASD **(Fig. 4a**, **Extended Data Fig. 7**, **Supplementary Table 6**). These include GeneM8, a microglial module containing genes involved in immune signaling and phagocytosis; and GeneM7, an immune response module containing genes such as NF-kB and interferon response pathways (**Fig. 4b**, **Supplementary Table 5-6**). Although neuronal and oligodendrocyte downregulation along with immune and microglia upregulation have been previously reported in ASD,^1,4–6^ these findings indicate that this dysregulation is widespread across the cerebral cortex, with increased magnitude in posterior regions, a pattern most pronounced in BA17.

**Figure 4.**
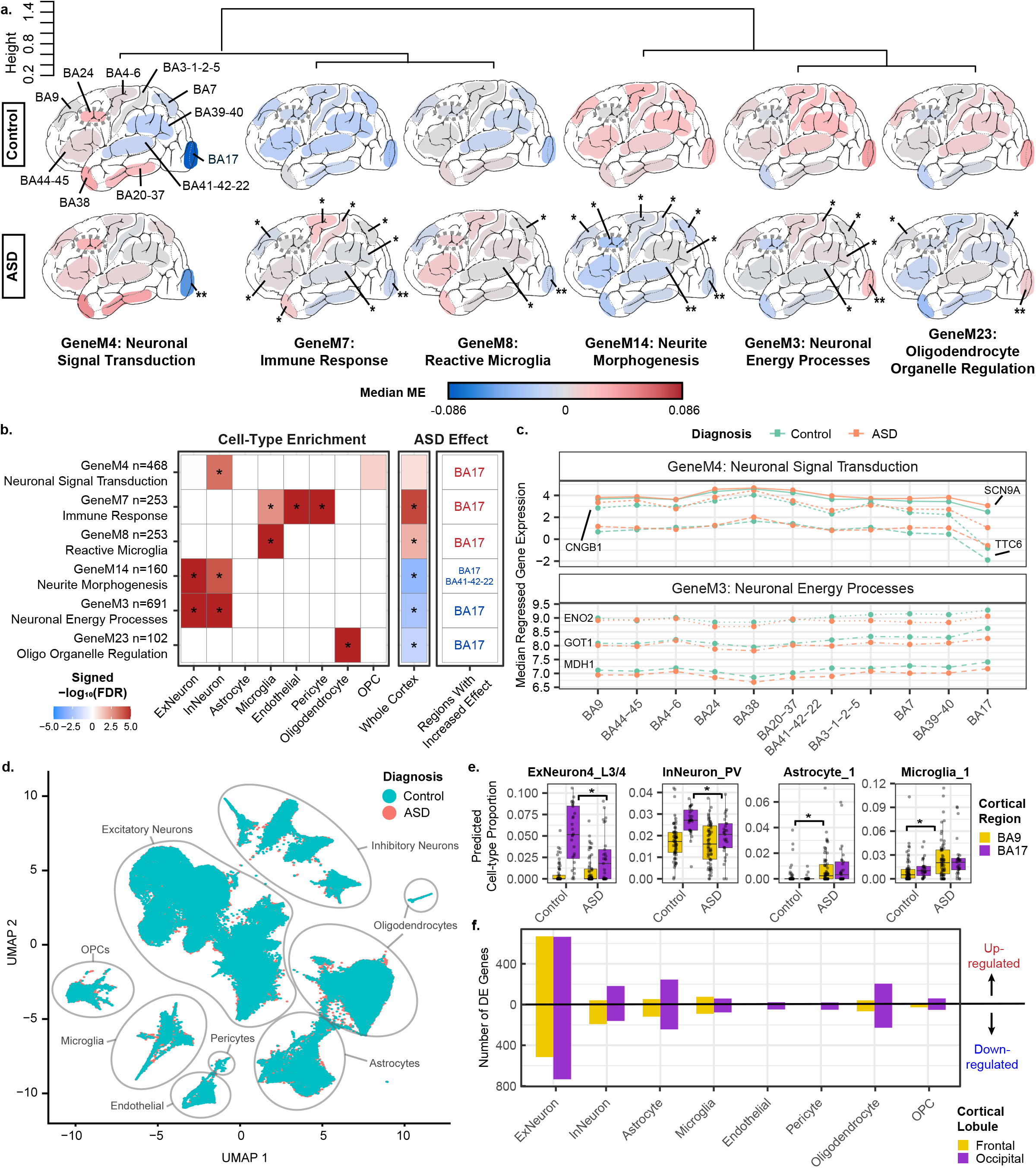
Functional Characterization of Regionally-variable Transcriptomic Dysregulation in ASD. **a.** Average linkage hierarchical clustering of the biweight midcorrelation of the top 6 most dysregulated gene co-expression module eigengenes (ME; first principal component of the module) with regionally-variable patterns of dysregulation. The median of the ME, stratified by diagnosis, is depicted for each cortical region examined. Significant region-specific dysregulation in ASD is marked with ‘*’, and regions with a significantly increased magnitude of effect compared to the whole cortex effect is marked with ‘**’ (**Methods**). **b.** Left and center, signed −log10(FDR) for cell-type enrichment (left) and whole cortex ASD effect for the modules depicted in **a** (center). ‘*’ indicates a significant enrichment (FDR < 0.05). Right, regions marked with ‘**’ in **a** are listed. Red text indicates upregulation and blue text indicates downregulation in ASD. ‘n’ is the number of genes in each module. **c.** Median regressed gene expression for the top three hub genes for GeneM4 (top) and GeneM3 (bottom). **d.** UMAP plot for snRNA-seq, containing matched ASD and neurotypical control samples from both the frontal (BA9 and BA4_6; ASD: 4, Control: 2) and occipital (BA17; ASD:2, Control: 2) cortical lobules. **e.** Predicted neural cell-type proportions obtained from cell-type deconvolution with bulk RNA-seq samples, using snRNA-seq cell-type markers. Cell-types with significant proportion differences in ASD are shown, with significant ASD changes marked with ‘*’. **f**. DE genes identified with snRNA-seq. Total number of down-regulated (bottom) and up-regulated (top) genes in ASD samples compared to controls is shown (the sum of all cell subtype DE genes within each broad cell-type).

The last three modules were observed to be significantly upregulated in ASD only in BA17. One of these, GeneM4, is an inhibitory neuron module containing many genes important for various intracellular signaling and maturation processes, such as *SCN9A* (**Fig. 4a-c**, **Supplementary Table 5-6**). Additionally, GeneM4 is significantly enriched for lincRNAs and for previously reported gene modules associated with upregulated pathways related to developmen^t5^ and signaling^1,6^ in ASD, although we observe this effect in BA17 for the first time (**Extended Data Fig. 7**). We also identified four other modules exhibiting strong region-specific dysregulation in regions other than BA17 (**Extended Data Fig. 7**). For example, the module GeneM34, which contains genes involved in cellular stress response regulatory processes, is upregulated with the greatest magnitude in BA4-6 and shows no significant effect in BA17 (**Extended Data Fig. 7**, **Supplementary Table 5-6**). None of the gene modules with regionally-variable magnitudes of ASD effect were significantly enriched for known ASD genetic risk variants.

#### Cell-type changes mirror regional variation

We finally sought to determine what might be driving the observed changes in magnitude of ASD effect across regions. It is well established that BA17 is the most neuronally dense region in the human brain, with a notable expansion in the thickness of L3/4, compared with other cortical regions.^18^ Likewise, there is an anterior-posterior gradient in neuronal density observed in mice and primates.^14–17^ As such, we posited that regional variation in cell density could be contributing to regional differences in magnitude of ASD effect. Regional neuronal density across multiple brain regions has not been quantitatively studied in the human brain, but such gradients have been established across some regions in non-human primates.^15,16^ Therefore, we compared the region-specific ASD effect size changes in our gene modules to regional neuronal nuclei density measured in primates^15^ for 6 matched regions across species. We observed a significant association between neuronal density and the effect sizes for several modules dysregulated in ASD (seven with FDR < 0.05, and an additional eight with FDR < 0.1, **Extended Data Fig. 9**, **Supplementary Table 7**). Further, L3/4 thickness was also associated with the region-specific ASD effect sizes in dysregulated modules (**Supplementary Table 7**).

These observations motivated us to perform single-nucleus RNA sequencing (snRNA-seq) in a small cohort of individuals to help evaluate how distinct neural cell-types could be contributing to the regional variance in ASD transcriptomic dysregulation identified with bulk RNA-seq (**Fig. 4d**, **Extended Data Fig. 9**, **Supplementary Table 7, Methods**). We sequenced over 150,000 nuclei from ASD and control samples across frontal and occipital cortices with matching bulk RNA-seq. From these data, we identified 35 distinct cell clusters and 4,953 cell-type-specific DE genes in ASD subjects in the frontal and occipital cortex. The vast majority of these were DE in excitatory neurons in both regions, and exhibited larger effects overall in the occipital lobe (**Fig. 4f**). While statistical power limited our ability to detect significant cell-type proportion differences between regions or diagnoses (**Methods**), we do observe that excitatory neurons are increased in proportion by ~5% in BA17 across both control and ASD subjects compared to frontal regions (**Extended Data Fig. 9**), corresponding with the primate neuronal density measurements. To predict how cell-type proportions may vary across our entire bulk RNA-seq dataset, next we utilized cell-type markers from our snRNA-seq to perform cell-type deconvolution in all samples (**Methods**). We identified 11 significant cell subtype proportion changes present across six different regions in ASD, characterized by neuronal decreases and astrocyte and microglia increases (**Fig. 4e**, **Extended Data Fig. 9**, **Supplementary Table 7**). We also found many anterior-posterior cell-type proportion gradients in control subjects that are attenuated in ASD (**Extended Data Fig. 9**, **Supplementary Table 7**), mirroring patterns observed with our bulk RNA-seq transcriptomic regional identity analysis.

When directly comparing cell-type-specific DE and deconvolved proportional changes in ASD with regional variability in the larger bulk transcriptome sample, we observed a convergent signal within excitatory neurons – in particular, those in L3/4 (ExNeuron4; **Fig. 4e-f**, **Extended Data Fig. 9**). Recapitulating the known increase in thickness of L3/4 in BA17 compared with other cortical regions^18^, we observed a significant increase in the estimated proportion of the ExNeuron4_L3/4 cluster in posterior regions, peaking in BA17 (**Fig. 4e; Extended Data Fig. 9g).**This regional pattern was significantly attenuated in ASD, with a ~2-fold median reduction in estimated Ex4 neuronal proportion in BA17 compared with controls. This cell cluster, with marker genes *RORB*, *PCP4*, *CUX2*, *PHACTR2*, and *EYA4*, also exhibited substantially greater cell-type-specific DE in snRNA-seq profiling from BA17 compared with frontal cortex (90 vs 0 DE genes, respectively; **Extended Data Fig. 9d**). Similarly, BA17 shows a substantially greater upregulation of inhibitory neuron genes in the single cell data, consistent with the observed greater up-regulation of GeneM4 (inhibitory neuron) in BA17 (**Fig. 4f**). Substantial changes in gene expression are also evident in other cell subtypes (**Extended Data Fig. 9, Supplementary Table 7**), such as microglia_2, which shows a strong and specific increase in DE genes in ASD BA17 compared to frontal regions. These observed intracellular/cell-type changes in neuronal and microglial gene expression are further supported by another snRNA-seq dataset containing a small ASD cohort, which assessed a single region.^19^ Here, through performing multi-region snRNA-seq and cell-type deconvolution, we show that predicted cell-type proportions as well as cell-type-specific gene expression profiles are impacted across the ASD cerebral cortex. Importantly, we see increased cell-type-specific transcriptomic dysregulation and lowered neuronal proportions with a notable convergence within L3/4 excitatory neurons in ASD BA17, a region where neuronal proportions are neurotypically abundant. These changes likely contribute to the pronounced ASD effect we observe with bulk RNA-seq in this region.

## Discussion

Overall, the findings presented here substantially expand our understanding of ASD pathology beyond the previously established ‘downregulated neuron’ and ‘upregulated glia/immune’ functional categories observed in frontal and temporal lobe. We identify gene and isoform expression changes in ASD that extend across the cerebral cortex, many neural cell-types, and specific biological processes (**Extended Data Fig. 10**), including primary sensory areas in addition to association areas.^1,4–6^ We find that the recently observed reactive astrocyte upregulation and blood-brain barrier membrane transport downregulation^1^ is extended cortex-wide in ASD. It is interesting to speculate that the substantial changes observed in area 17, a primary sensory region, may be related to the widespread observation of sensory hypersensitivity or processing abnormalities in ASD.^40^ Nevertheless, this region shows the most profound changes in gene expression, a clear demonstration that these alterations are not specific to higher association areas.

Furthermore, we find that other dysregulated pathways observed before in ASD - particularly upregulated immune response and reactive microglia genes, along with downregulated neurite morphogenesis and neuronal energy pathway genes - are not only impacted cortex-wide in ASD, but impacted in a regional gradient that reflects fundamental elements of cortical cytoarchitecture, such as neuronal density. It is also notable that the magnitude of region-level differences in ASD parallels regional variance in attenuation of transcriptomic identity (ARI gene dysregulation), suggesting that they reflect related processes. That the gradient of region-specific changes between ASD and controls coincides with both neuronal proportion differences and cell-type-specific transcriptomic dysregulation further suggests that the interplay of cytoarchitecture and cell-type gene expression, rather than a single one of these features, influences our ability to observe transcriptomic changes in bulk tissue. Given the connection between regional cytoarchitecture, local circuits and long-range brain connectivity,^20,21^ parsimony suggests that in addition to developmental patterning contributions,^5,22^ the diminution of transcriptomic regional identity reflects changes in local neuronal circuit dysfunction and deficits in synaptic plasticity and homeostasis that are widely propagated.^20^ This is supported by our observation that the gene co-expression module representing synaptic plasticity genes is downregulated cortex-wide and is significantly enriched with common and rare ASD genetic risk variants, further emphasizing that synaptic plasticity is a convergent pathway in ASD. Given this result, along with our observations of profound neuronal dysregulation present throughout the ASD cortex, future work should determine which specific aspects of synaptic plasticity may contribute to causal mechanisms in the disorder across specific brain regions and developmental timepoints.

Several additional factors should guide the interpretation of these results. The samples utilized in this work were obtained from heterogeneous postmortem cortical tissue, meaning that the results reported here are broadly applicable to the postnatal ASD cortex across both sexes and a span of ages from two to 68 years old, and they should be interpreted in this context. Rigorous methodology was utilized at every step to account for biological and technical variability, ensuring that the results reported here are conservative and widely applicable. Additionally, bulk tissue RNA-seq, in contrast to single cell and nucleus RNA-seq, does not have the cellular resolution to assess dissection variability across cortical regions and cell-type specificity of transcriptomic changes. We addressed this by performing snRNA-seq, which significantly enhanced our understanding of regional variation in ASD transcriptomic dysregulation. However, snRNA-seq also has its own limitations. While snRNA-seq can profile tens of thousands of cells, snRNA-seq experiments typically have fewer unique samples than bulk RNA-seq experiments, and the comparability of snRNA-seq cell-type proportions to true sample cell-type proportions is currently unclear.^23^ It is also challenging to estimate isoform quantifications with single cell RNA-seq approaches, whereas this remains a strength of bulk tissue RNA-seq.^24^ Leveraging this, we subsequently identified an upregulated isoform-specific co-expression module enriched with ASD GWAS variants, implicating increased protein folding dysfunction for the first time as a putative pathway contributing to ASD causal mechanisms. Interestingly, upregulated proteostasis is also implicated in Down’s Syndrome,^25,26^ supporting that protein folding machinery may be an affected biological process in multiple neurodevelopmental disorders. The utilization of methods that have greater cellular resolution is necessary for the improved and continued mapping of the results presented here to specific cortical cell-types. As we seek to gain a complete understanding of ASD neural pathology, future approaches which integrate different sources of biological data - including this cortex-wide transcriptomic resource - to determine how ASD risk genes are acting in the brain will be essential.

## Methods

### Sample Acquisition and Preparation for RNA-seq

Postmortem cortical brain samples were acquired from the Harvard Brain Bank as part of the Autism BrainNet project (formerly the Autism Tissue Project, ATP) and the University of Maryland Brain Banks (UMDB). A total of 842 samples from subjects with ASD, dup15q syndrome, and non-psychiatric controls (112 unique subjects) across 11 cortical regions encompassing all major cortical lobes – frontal: BA4/6, BA9, BA44/45, BA24; temporal: BA38, BA41/42/22, BA20/37; parietal: BA3/1/2/5, BA7, BA39/40; and occipital, BA17 - were acquired. These included 253 samples previously published in Parikshak et al., Nature 2016^5^ from BA9 and BA41/42/22 and/or Gandal et al., Science 2018b^1,5^ from BA9, BA4/6, and BA41/42/22. An ASD diagnosis was confirmed by the Autism Diagnostic Interview-Revised (ADIR) in 30 of the subjects. In the remaining 19 subjects, diagnosis was supported by clinical history. Frozen brain samples were stored at −80 deg C. To extract RNA from these samples, first approximately 50-100mg of tissue were dissected from the cortical regions of interest on dry ice in a dehydrated dissection chamber to reduce degradation effects from sample thawing or humidity. Then, RNA was isolated from each sample using the miRNeasy kit with no modifications (Qiagen). For each RNA sample, RNA quality was quantified using the RNA Integrity Number (RIN) on an Agilent Bioanalyzer.

### RNA-seq and RNA Data Processing

Initial sequencing in BA9 and BA41/42/22 was performed in three batches as published by Parikshak et al., Nature 2016.^5^ The remaining regions, along with additional BA9 and BA41/42/22 samples, were sequenced across three new batches. For all of these batches, strand-specific RNA-seq libraries were prepared. For the first two batches, the TruSeq Stranded Total RNA sample prep kit with RiboZero Gold (Illumina) was used to obtain rRNA-depleted libraries. The remaining batch was prepared with the TruSeq RNA Exome sample prep kit (formerly the TruSeq RNA Access sample prep kit; Illumina). All libraries were randomly pooled to multiplex 24 samples per lane using Illumina TruSeq barcodes. Each lane was sequenced five times on an Illumina HiSeq 2500 or 4000 instrument using high output mode with standard chemistry and protocols for 50, 69, or 100 bp paired-end reads (read length varied by batch) to achieve a target depth of 70 million reads.

After sequencing, the resulting sample FASTQ files from all batches (including the Parikshak et al.^5^ samples) were subjected to the same processing pipeline. First, FASTQ files were assessed with FastQC^27^ (v0.11.2) to verify that quality was sufficient for further processing. FASTQ files were then aligned to the human reference genome (GRCh37^28^ Ensembl v75) with STAR^29^ (v2.5.2b). Picard tools^30^ (v2.5.0) was used with the resulting BAM files to collect various read quality measures, in addition to the quality measures collected by STAR. verifyBAMID^31^ was also used with these BAM files along with known sample genotypes from Parikshak et al.^5^ to validate that sample identity was correct for all BAM files. Additionally, the expression of XIST (a female-specific gene) was assessed to contribute to sample identity verification. Finally, RSEM^32^ (v1.3.0) was used for quantification (Gencode^33^ release 25lift37) to obtain expected read counts at the gene and isoform levels.

Expected gene and isoform read counts were then subjected to several processing steps in preparation for downstream analysis, mainly using R.^34^ First, Counts Per Million (CPM) were obtained from counts for gene and isoform filtering purposes. Genes and isoforms were filtered such that genes/isoforms with a CPM > 0.1 in at least 30% of samples were retained. Genes/isoforms were also removed which had an effective length (measured by RSEM) of less than 15 bp. Isoforms were additionally filtered such that all isoforms corresponded with genes in the gene-level analysis. The counts for the remaining genes (24,836) and isoforms (99,819) passing these filters were normalized using the limma-trend approach in the limma^35^ R package.

Briefly, the limma-trend approach obtains normalized expression data through taking the log2(CPM) of read counts with an adjustment for sample read depth variance. An offset value calculated with CQN^36^ accounting for GC content bias and gene/isoform effective length bias in read quantification was also incorporated during the normalization process. With this normalized expression data, sample outliers were identified in each sequencing batch by cortical lobe (frontal, parietal, temporal, and occipital) group that had both (1) an absolute z-score greater than 3 for any of the top 10 expression principal components (PCs) and (2) a sample connectivity score less than −2. Sample connectivity was calculated using the fundamentalNetworkConcepts function in the WGCNA^10^ R package, with the signed adjacency matrix (soft power of 2) of the sample biweight midcorrelation as input. This process identified 34 outliers, resulting in a final total of 808 samples (341=Control, 384=ASD, 83=dup15q) which were carried forward for analysis.

### Evaluating Previous Co-Expression Modules and ASD DE Genes/Isoforms Cortex-wide

#### Linear models for all subsequent analyses are described in the Supplementary Methods

To determine how gene co-expression modules previously identified in Parikshak et al.^5,35^ and Voineagu et al.^4^ were effected across distinct cortical regions, we first created a regressed gene expression dataset that only contained the effects of biological covariates (subject, diagnosis, region, sequencing batch, sex, ancestry, age, and age^2^). This regressed dataset was created with the ‘lmerTest’^37^ package in R through subtracting the effects of technical covariates from each gene, leaving only the random intercept, biological covariate effects, and the residual. ASD-associated module eigengene region-specific ASD effects were identified using contrasts (eg. Control_BA17 - ASD_BA17) with the limma^34^ R package with this regressed expression dataset, accounting for all biological covariates. Region-specific contrasts with a p-value < 0.05 were considered significant (FDR-correction was unwarranted since only eight module eigengenes were examined).

To identify genes and isoforms dysregulated in ASD both within specific regions and cortex-wide, the limma^35^ R package was applied with the gene and isoform expression data using our full gene and isoform models (both biological and technical covariates). The standard limma^35^ workflow was implemented as recommended for linear mixed models. Region-specific dysregulation was identified as described above for the Parikshak et al.^5^ and Voineagu et al.^4^ modules. Whole cortex dysregulation was established through subtracting the sum of the ASD region-specific effects from the sum of the Control region-specific effects. For both region-specific and whole cortex effects, genes and isoforms with an FDR-corrected p-value < 0.05 were considered significantly dysregulated. dup15q region-specific and whole cortex dysregulation was also established in this manner. The fixed effects of sex, age, and age^2^ were also acquired (shared in **Supplementary Table 3**) using the full gene and isoform models.

The methodology used to evaluate region-specific ASD effects compared to whole cortex ASD effects is described in the Supplementary Methods.

### Transcriptomic Regional Identity Analysis

To identify differentially expressed genes and isoforms between all 55 pairs of cortical regions, a regressed gene expression dataset containing only the random effect of subject and the fixed effects of diagnosis and region (along with the model residual) was used. Regression was performed as described for evaluation of previously identified co-expression modules. Significant attenuation of DE genes between each pair of regions (a reduction in transcriptomic regional identity differences) in ASD was established through the following process. (1) ASD and Control subjects containing each region in the regional pair were extracted for use in the analysis. (2) Separately in ASD and Control subjects, the number of DE genes between regions was calculated using the paired Wilcoxon signed-rank test. Genes with an FDR-corrected p-value < 0.05 were considered DE. (3) The difference in the number of DE genes between regions for ASD v Control subjects was calculated (the ‘true’ difference). (4) A permuted distribution of the difference in DE genes between regions for ASD v Control subjects was generated to test the ‘true’ difference. Each permutation (10,000 in total) randomly assigned ‘ASD’ and ‘Control’ status to subjects, but kept the number of ASD and Control subjects consistent with the true number of ASD and Control subjects. (5) A two-tailed p-value was obtained from testing the ‘true’ difference against the permuted distribution. If the regional comparison p-value < 0.05, with the number of DE genes between regions in ASD less than that in Controls, then the regional comparison was considered significantly attenuated in ASD. Otherwise, the regional comparison was considered over-patterned in ASD. This procedure was repeated with isoform level regressed gene expression data (similarly, only containing the random effect of subject and the fixed effects of diagnosis and region, along with the model residual) to identify altered transcriptomic identities in ASD at the isoform-level.

The previously described permutation approach was designed to identify differences in transcriptomic regional identity in ASD. Importantly, this method is not appropriate for assessing variance in expected numbers of DE genes between regions across regional pairs and diagnoses, since the number of ASD and Control subjects varied across regional pairs. To examine this, for each regional comparison we subset to 10 pairs of ASD and Control subjects (10 was selected since every regional comparison had at least this many subjects). When subsetting, subjects were removed such that the remaining subjects were closest in age to the median age of the available samples for that regional comparison. A bootstrap approach was then used to calculate the number of DE genes (p-value < 0.05) between regions separately in Control and ASD subjects through sampling subjects with replacement (mean taken across 10,000 bootstraps). The same regressed expression dataset used for the permutation approach was utilized for this bootstrap analysis. Any regional comparison in which the number of DE genes between regions was less in ASD than in Control subjects was considered trending towards attenuation in ASD.

To validate our bootstrapped estimates for the number of DE genes between pairs of regions in Controls, we compared these estimates to those of the Allen Brain Atlas^9^, which is the best publicly available work for comparison. Allen Brain Atlas regions were matched to Brodmann regions (**Supplementary Table 4**) and matching regional pairs were extracted for comparison with this work. When the Allen Brain Atlas had two or more regional pairs matching one regional pair in this work, the mean was taken across the Allen Brain Atlas regional pairs. A p-value for the association of the number of DE genes between regions in Controls obtained in this work compared to the Allen Brain Atlas was calculated from a linear model (cortex-wide bootstrap mean ~ allen brain atlas mean).

We applied a stringent filtering process to identify high-confidence attenuated regional identity (ARI) genes from each significantly attenuated regional comparison identified with the permutation procedure described above. First, for each of the attenuated regional comparisons, we extracted the genes which were identified as DE between regions in subjects labeled as Controls in each of the 10,000 permutations. Then, we calculated how many times each of the genes truly DE between pairs of regions in the Control subjects were present in their respective permuted groups (ranging from a possible 0 to 10,000 occurrences). Those ‘true’ DE genes which were present in less than 95% of their respective permutations were retained as ARI genes for each attenuated regional comparison. For each set of ARI genes (ten total), each gene was matched to the region in which it had higher expression in Control subjects. The paired Wilcoxon signed-rank p-values identified for these genes in Controls (those subjects used for the permutation analysis) were also extracted and are shared in **Supplementary Table 4**.

ARI gene groups (ARI downregulated genes, those highly expressed in BA17 and BA39-40 relative to other regions in Controls; ARI upregulated genes, those lowly expressed in BA17 and BA39-40 relative to other regions in Controls) were created through taking the union (without duplicates) across all ten identified ASD-attenuated regional comparisons, and sorting genes into the two groups based on gene expression profiles across regions. The details of this process are described in the Supplementary Methods, along with functional annotation procedures.

### Network-Based Functional Characterization

Standard workflows, as previously described in Parikshak et al.^5^ and Gandal et al.^,1^ were followed (with minor modifications) to identify gene and isoform co-expression modules using Weighted Gene Correlation Network Analysis (WGCNA).^10^ Details regarding network formation, module identification, and module functional characterization are described in the Supplementary Methods.

### snRNA-seq and Cell-type Deconvolution

Cell types were annotated based on expression of known marker genes visualized on the UMAP plot, violin plots, and by performing unbiased gene marker analysis. To gain insight into the regional enrichment or diagnostic enrichment of cell types, the relative proportion of the number of nuclei in each cell type was normalized to the total number of nuclei captured from each library. Average cell-type proportions and standard errors (across libraries) were scaled such that each Lobule x Diagnosis group sums to 100%, so that cell-type proportions in these groups could be fairly compared across all cell-types. To determine if any changes in cell-type proportion were statistically significant, we implemented scDC^38^ to bootstrap proportion estimates for our samples (**Supplementary Table 7**). We employed a linear mixed model (random effect of subject) to determine if any changes in cell-type proportion were present across regions and diagnoses. None of the model covariates were statistically significant (p > 0.05 for all model covariates). However, we did find several significantly different predicted cell-type proportions in ASD with cell-type deconvolution analysis. We describe methods for cell-type deconvolution in detail in the Supplementary Methods. To identify genes differentially expressed in ASD compared to control in each cell type, the non-parametric Wilcoxon rank sum test was applied including gene detection rate and sequencing depth within the model. We compared frontal cortex ASD cells to frontal cortex control cells within each cluster and likewise for the occipital cortical cells. The bars in **Figure 4e** are the summation of all differentially expressed genes identified in each cell subtype for the broader cell-type (eg. all excitatory neuron subtype DE genes are summed to obtain the number of DE genes in the broad excitatory neuron cell class). Further details regarding the snRNA-seq analysis are included in the Supplementary Methods.

## Supporting information

Supplementary Methods, Supplementary Tables, and Supplementary References

## Data availability

All of the raw bulk RNA-seq data (FASTQ files) and processed (utilized for DE gene analysis, transcriptomic regional identity analysis, and WGCNA) bulk RNA-seq data that support the findings of this study will be deposited in a publicly accessible repository. snRNA-seq data will be made available by the corresponding authors upon reasonable request. All of the code, raw data, and processed data for the bulk RNA-seq analysis that support the findings of this study will also be made available in a publicly accessible GitHub repository, where readers may also access an R Shiny tool to visualize RNA-seq data across the biological covariates assessed in this study.

## Code availability

All of the code, raw data, and processed data for the bulk RNA-seq analysis that support the findings of this study will also be made available in a publicly accessible GitHub repository, where readers may also access an R Shiny tool to visualize RNA-seq data across the biological covariates assessed in this study.

## Extended Data Figure Legends

**Extended Data Figure 1 | Experiment Workflow and Sample Overview.** a. Overview of experiment workflow. b. Summary of sample composition (biological data, brain bank source, and PMI).

**Extended Data Figure 2 | Quality Control Measures.** a. Sequencing batch parameters. b. Sequencing batches (top) and sequencing statistics (bottom) by region and diagnosis. c. Top 15 expression PCs (gene and isoform, with % of variance explained denoted) association with meta data (top) and sequencing statistics (bottom).

**Extended Data Figure 3 | Model Covariates and Previous Studies Across 11 Cortical Regions.** a. For the covariates selected for the gene (left) and isoform (right) linear mixed models, % of expression variance explained across all genes/isoforms. b-c. For the Voineagu et al. and Parikshak et al. studies, b. ASD associated gene module ASD effect (standard error bars and cortical lobes indicated) and c. ASD log2 FC of DE genes identified in these studies, compared to this dataset (Spearman’s correlation rho, R, is plotted along with the linear least squares regression best fit line).

**Extended Data Figure 4 | Transcriptomic Changes Across 11 Cortical Regions.** a. Overlap of Whole-Cortex DE ASD genes and isoforms (blue) with other cortical region DE genes (no color). Regions with no third numeric label on the right completely overlap with the Whole-Cortex DE genes. b. For the Whole-Cortex DE, overlap of genes and isoforms. Regions not shown have no unique DE. c. log2(FC) (top) and standard error (SE, bottom) of the Whole-Cortex ASD DE overlapping and distinct genes and isoforms. d. Overlap in DE ASD and dup15q genes and isoforms. e. For regions with DE ASD genes (left) and isoforms (right), ASD log2(FC) v. dup15q log2(FC) for specific regions (with principal components regression slope, S).

**Extended Data Figure 5 | Transcriptomic Regional Identity Attenuation in ASD.** a. Mean-centered distribution of 10,000 permutations for the significantly attenuated regional comparisons in ASD. Red bar = true difference in the number of DE genes between controls and ASD. b. Sample size for all regional comparisons. Permutation p-values for all regional comparisons. d. For 10,000 regional comparison bootstraps, ratio of DE genes in ASD compared to controls. e. Number of DE genes between pairs of regions in this study (mean across bootstraps in controls, y-axis) compared to the Allen Brain Atlas (ref. 10, mean across matched regions, x-axis; see Methods for matched regions). This Allen Brain Atlas dataset, with only 2 unique brains, is the best publicly available dataset for comparison (linear least squares regression best fit line plotted).

**Extended Data Figure 6 | Additional ARI gene dysregulation.** a. First principal component (PC1) of posteriorly downregulated (1,881, left) and upregulated (1,695, right) ARI genes identified in ASD, plotted in Controls and dup15q (loess regression line plotted). b. For each significantly attenuated regional comparison, the identified attenuated regional identity (ARI) genes. At center, number of ARI genes with greater neurotypical expression in each pair of regions. On either side of the barplot, the PC1 of the genes with greater neurotypical anterior (left) or posterior (right) expression is plotted across the pair of regions in Controls and ASD. The Wilcoxon signed-rank test (unpaired) p-value is shown.

**Extended Data Figure 7 | Gene-level Co-Expression Network Analysis Module Associations.** Top: average-linkage hierarchical clustering of module eigengene biweight midcorrelations. Significant FDR corrected p-values are indicated (FDR < 0.05; for GWAS, FDR < 0.1). Any signed −log10(p) colors greater or less than 5/-5 are set at a max/min of 5/-5. For ASD, dup15q, and Age covariates, FDR p-value from the linear mixed model testing the association of these covariates with module eigengenes is depicted. For the ASD and dup15q region-specific comparisons, cortical lobule colors are indicated (Fig. 1a), and bold-italic FDR p-values indicate that these regions are effected with significantly greater magnitude than the ASD whole-cortex (Methods). For gene biotypes, both positive and negative enrichment is shown (Methods). Positive enrichment is shown for cell-type, neuronal subtype (ref: Hodge et al, Nature 2019), ARI gene, GWAS, and rare variant enrichment (Methods).

**Extended Data Figure 8 | Isoform-level Co-Expression Network Analysis Module Associations.** Top: average-linkage hierarchical clustering of module eigengene biweight midcorrelations. Significant FDR corrected p-values are indicated (FDR < 0.05; for GWAS, FDR < 0.1). Any signed −log10(p) colors greater or less than 5/-5 are set at a max/min of 5/-5. For ASD, dup15q, and Age covariates, FDR p-value from the linear mixed model testing the association of these covariates with module eigengenes is depicted. For the ASD and dup15q region-specific comparisons, cortical lobule colors are indicated (Fig. 1a). For gene biotypes, both positive and negative enrichment is shown (Methods). Positive enrichment is shown for cell-type, GWAS, and rare variant enrichments (Methods).

**Extended Data Figure 9 | Neuronal Density Associations, snRNA-seq, and Cell-type Deconvolution.** a. Macaque neuronal density v. module eigengene ASD effect for modules featured in Fig. 4c-d (linear least squares regression). Both p-value and FDR corrected p-value are plotted. b. P-value histogram of all gene modules’ linear least squares regression with macaque region-specific neuronal density. c. UMAP plots of snRNA-seq with cell sub-types (top) and brain regions (bottom) depicted. d. Number of genes differentially expressed in ASD in each cell subtype. Upregulated genes are above 0 (red arrow) and downregulated genes are below 0 (blue arrow). e. Average proportion of each broad cell-type in each diagnosis x cortical lobule group, derived directly from the snRNA-seq data. f. Additional significant (Bonferroni corrected p-value < 0.05) cell-type proportion differences in ASD from cell-type deconvolution. Region and cell-type are indicated in the title of each plot. g. For two example cell-types, cell-type proportion attenuation in ASD across regions. ANOVA p-values stratified by diagnosis are shown.

**Extended Data Figure 10 | Results Summary.** Overview of RNA-sequencing experiment and results. Region-specific dysregulation scale in the top right corner and the leftmost portion of the bottom panel depict the region-specific slopes compared to the whole cortex effect from Fig 1d. Median PC 1 of the ARI dysregulated genes is plotted in the middle panel. In the right portion of the bottom panel, the median ME of GeneM4 (left) and GeneM3 (right) is depicted.

## Acknowledgements

Tissue, biological specimens or data used in this research were obtained from the Autism BrainNet (formerly the Autism Tissue Program), which is sponsored by the Simons Foundation, and the University of Maryland Brain and Tissue Bank, which is a component of the NIH NeuroBioBank. We are grateful to the patients and families who participate in the tissue donation programs. Funding for this work was provided by grants to D.H.G (NIMH R01MH110927, U01MH115746, P50-MH106438, and R01 MH-109912, R01 MH094714), grants to MJG (SFARI Bridge to Independence Award, NIMH R01-MH121521, NIMH R01-MH123922, NICHD-P50-HD103557), grants to JRH (Achievement Rewards for College Scientists Foundation Los Angeles Founder Chapter, UCLA Neuroscience Interdepartmental Program). We also thank Janet Sinsheimer for discussion of the transcriptomic regional identity analysis methodology.

## Author Contributions

J.R.H contributed to the entire manuscript, including all analysis of the bulk RNA-seq data, supporting analysis of the snRNA-seq data, writing the main text, creating all figures, and compiling all supplementary information.

B.W. completed the snRNA-seq experiment with matched samples, and also performed the main cell-type proportion and DE gene analyses with the snRNA-seq data. B.W. also contributed to Figure 4.

G.T.C. completed an experiment with human *in vitro* neuronal cultures over-expressing ETV4, to determine if this gene may be a transcriptional regulator of some of the attenuated regional identity genes identified in this work. Results were inconclusive, and were ultimately not included in this manuscript.

S.P. completed all of the raw data processing of the bulk RNA-seq data (obtaining gene and isoform level quantifications and RNA-seq quality metrics).

P.E. conducted the cell-type deconvolution analysis.

N.C. contributed to the cell-type deconvolution analysis.

G.D.H provided expert functional circuit and regional neuroanatomy input for the manuscript and assisted with figure design.

D.A. carried out the protein-protein interaction enrichment analysis for gene and isoform modules.

G.K. assisted with making the GitHub repository and made the R Shiny data visualization application accompanying this work.

G.R. provided assistance with the bulk RNA-seq analysis data interpretation and visualization.

C.L.H. created the RNA-seq computational pipeline used to process the raw bulk RNA-seq data and obtain gene and isoform level gene expression quantification.

T.J. contributed to the cell-type deconvolution analysis.

D.W. contributed to the cell-type deconvolution analysis.

J.O. extracted RNA and prepared RNA libraries for the bulk RNA-seq.

Y.E.W. assisted with sample dissection for the bulk RNA-seq RNA extraction.

N.N.P. contributed to the initial planning of this experiment and assisted with sample dissections for the bulk RNA-seq RNA extraction.

V.S. contributed to the initial planning of this experiment and assisted with sample dissections for the bulk RNA-seq RNA extraction.

G.B. contributed to the initial planning of this experiment.

M.Ge. served in an advisory and project-planning role for the cell-type deconvolution analysis.

B.P. provided expert statistical advice for the bulk RNA-seq analysis, particularly for the permutation tests assessing region-specific increases in magnitude of ASD gene expression effects.

M.J.G. provided project leadership, expert advice, substantial contributions to the entire manuscript, and mentorship for J.R.H. along with funding for this work. M.J.G also contributed to sample dissections for the bulk RNA-seq RNA extraction.

D.H.G. provided project leadership, expert advice, substantial contributions to the entire manuscript, and mentorship for J.R.H. along with funding for this work. D.H.G also obtained all of the samples for RNA-seq.

