## Extended Data Figures for "Broad transcriptomic dysregulation across the cerebral cortex in ASD"

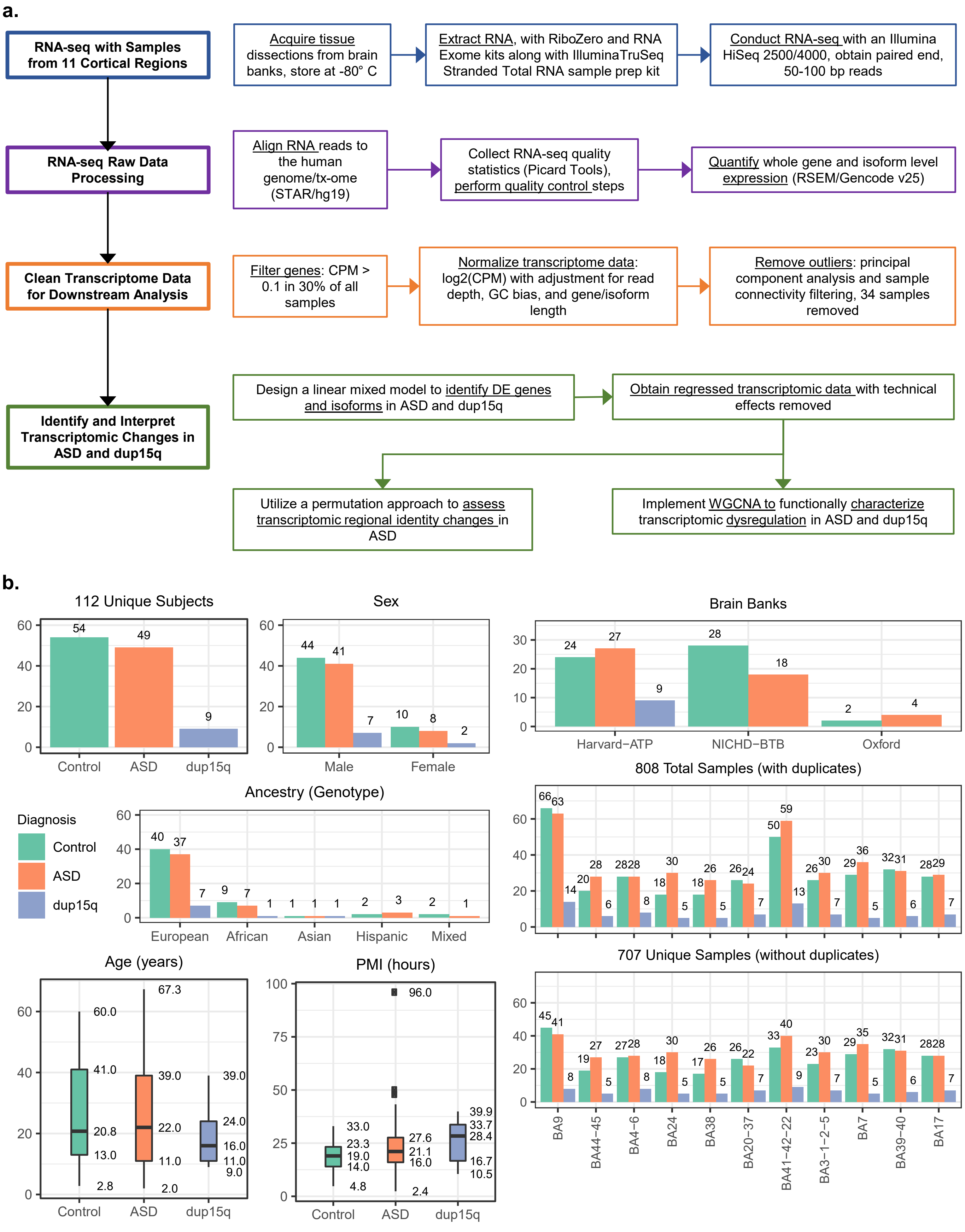


**Extended Data Figure 1 | Experiment Workflow and Sample Overview. a.** Overview of experiment workflow. **b.** Summary of sample composition (biological data, brain bank source, and PMI).

**a.**

| Batch Name | Batch Year | Sequencing Strand-Specificity | Read Length (paired end) |
|---|---|---|---|
| ASD3Reg_Batch1 | 2012 | Unstranded | 50 |
| ASD3Reg_Batch2 | 2013 | Unstranded | 50 |
| ASD3Reg_Batch3 | 2015 | Unstranded | 50 |
| ASDPan_Batch1 | 2015 | First-Strand | 50 |
| ASDPan_Batch2 | 2016 | First-Strand | 69 |
| ASDPan_Batch3 | 2015 | First-Strand | 100 |

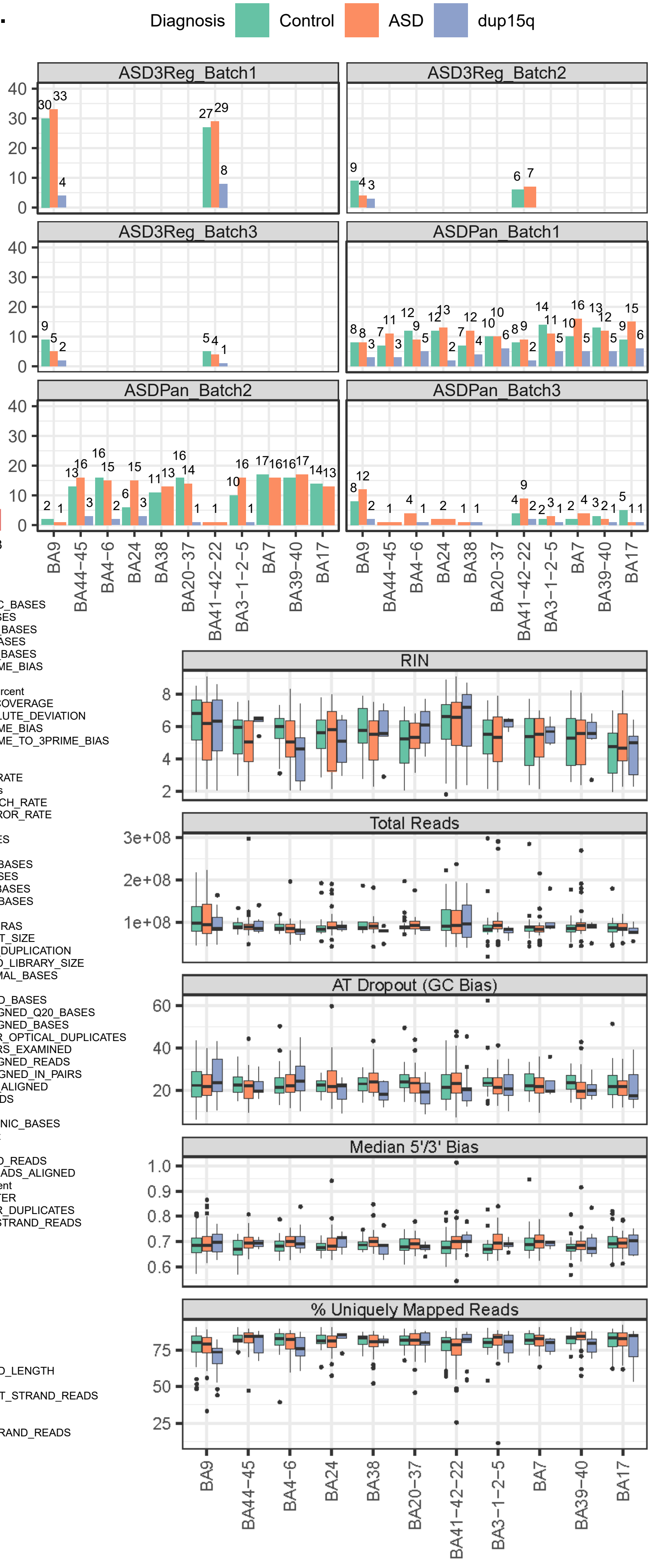


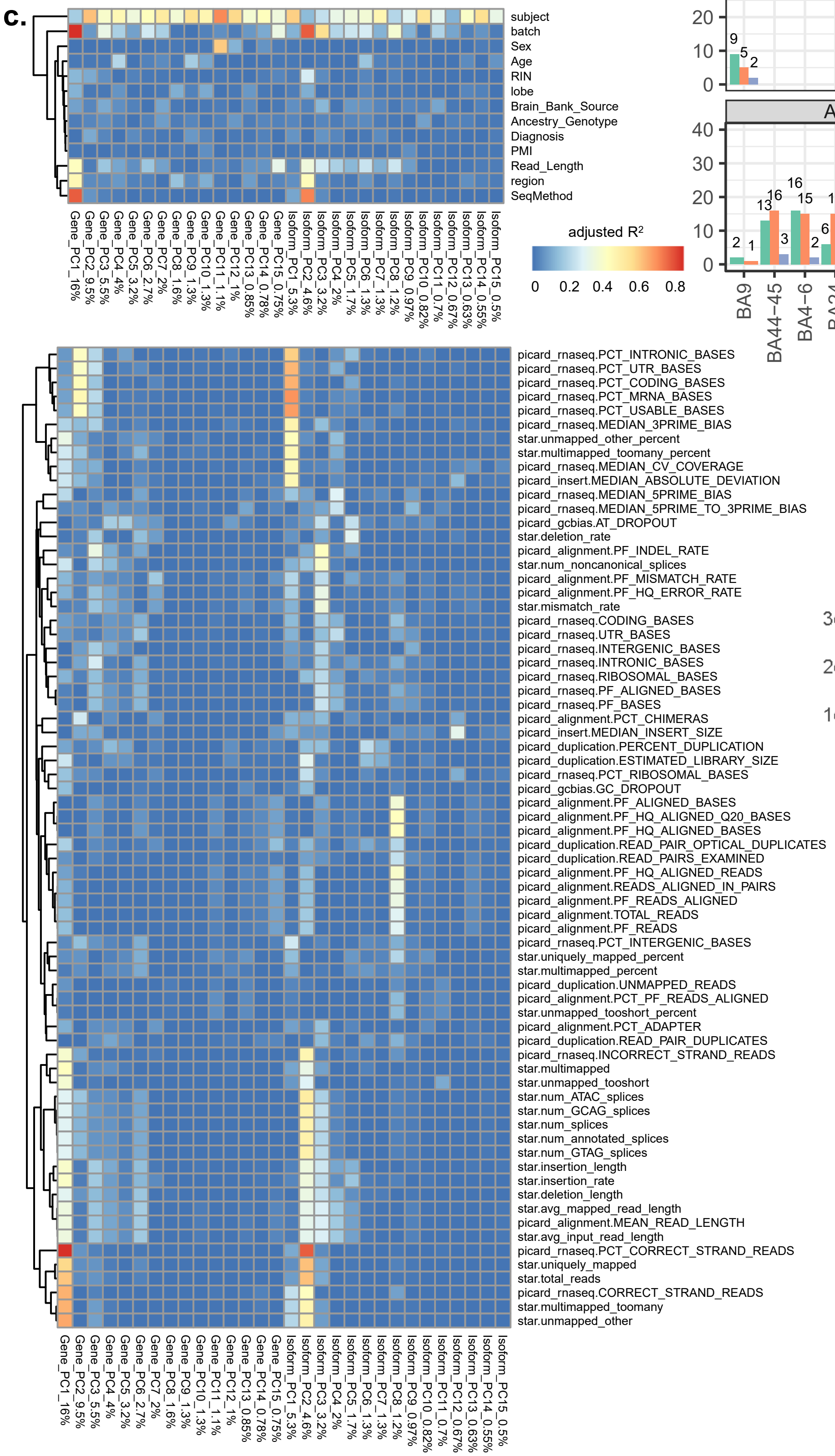


**Extended Data Figure 2 | Quality Control Measures. a.** Sequencing batch parameters. **b.** Sequencing batches (top) and sequencing statistics (bottom) by region and diagnosis. **c.** Top 15 expression PCs (gene and isoform, with % of variance explained denoted) association with meta data (top) and sequencing statistics (bottom).

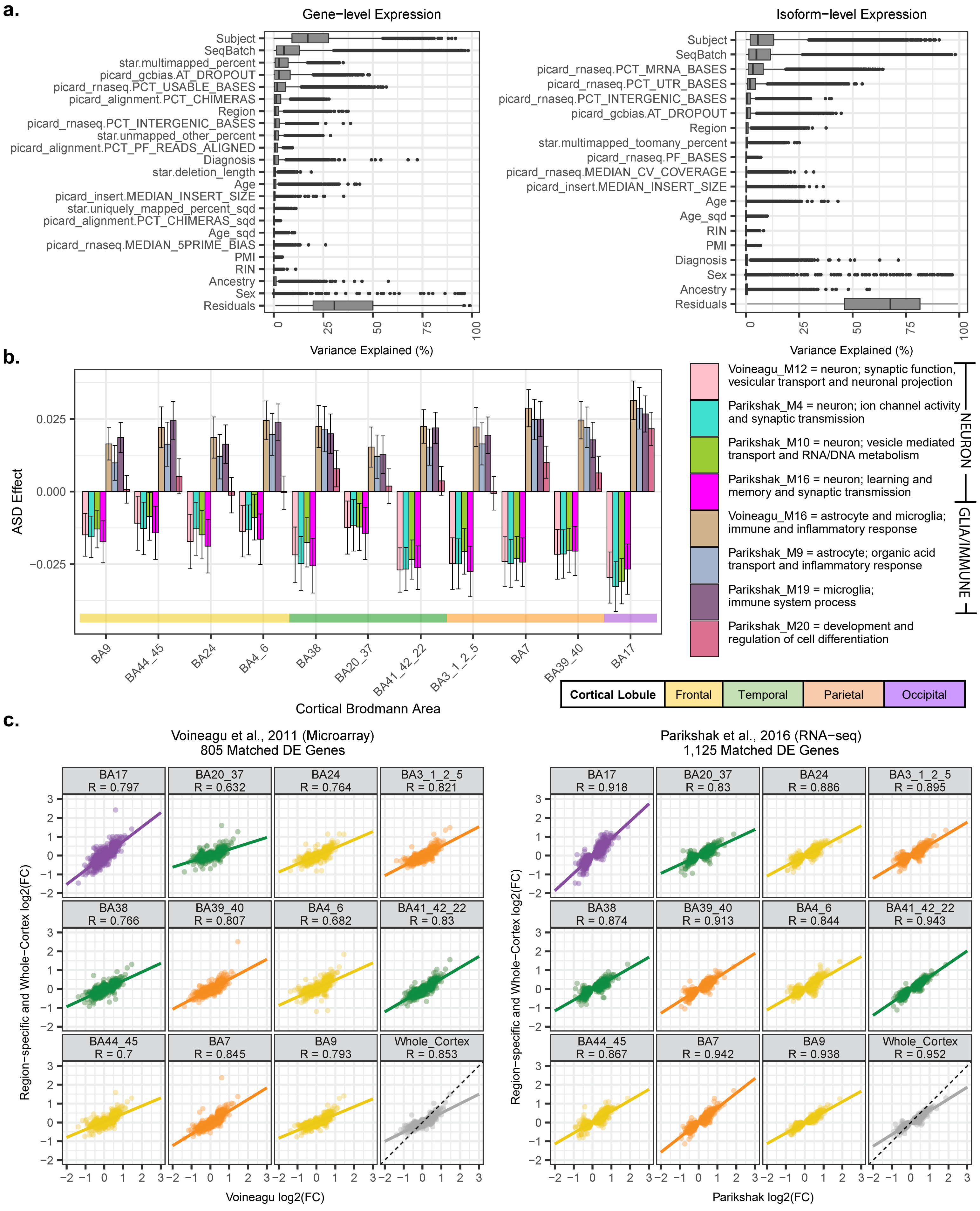


**Extended Data Figure 3 | Model Covariates and Previous Studies Across 11 Cortical Regions. a.** For the covariates selected for the gene (left) and isoform (right) linear mixed models, % of expression variance explained across all genes/isoforms. **b-c.** For the Voineagu et al. and Parikshak et al. studies, **b.** ASD-associated gene module ASD effect (standard error bars and cortical lobes indicated) and **c.** ASD log2 FC of DE genes identified in these studies, compared to this dataset (Spearman's correlation rho, R, is plotted along with the linear least squares regression best fit line).

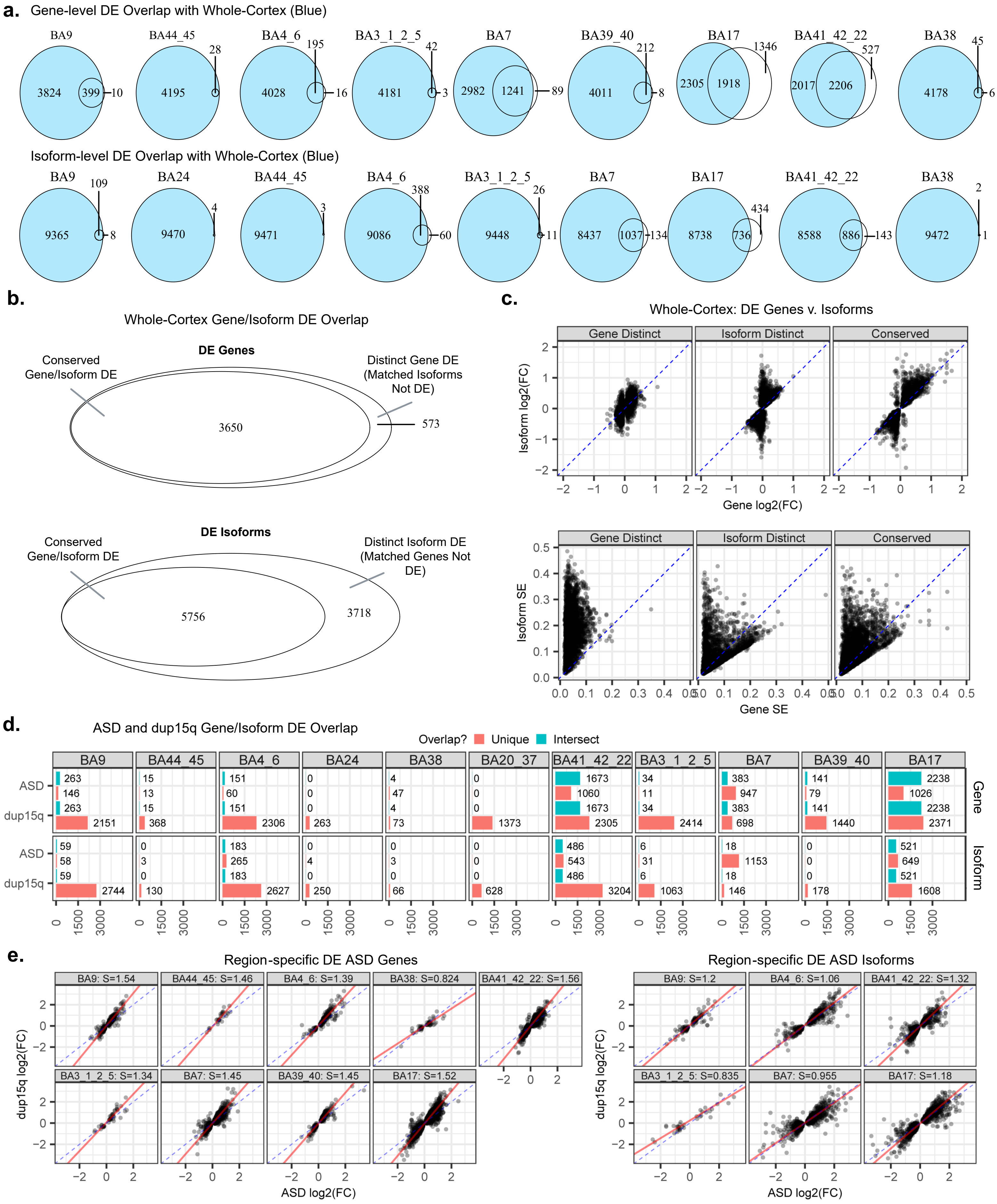


**Extended Data Figure 4 | Transcriptomic Changes Across 11 Cortical Regions. a.** Overlap of Whole-Cortex DE ASD genes and isoforms (blue) with other cortical region DE genes (no color). Regions with no third numeric label on the right completely overlap with the Whole-Cortex DE genes. **b.** For the Whole-Cortex DE, overlap of genes and isoforms. Regions not shown have no unique DE. **c.** log2(FC) (top) and standard error (SE, bottom) of the Whole-Cortex ASD DE overlapping and distinct genes and isoforms. **d.** Overlap in DE ASD and dup15q genes and isoforms. **e.** For regions with DE ASD genes (left) and isoforms (right), ASD log2(FC) v. dup15q log2(FC) for specific regions (with principal components regression slope, S).

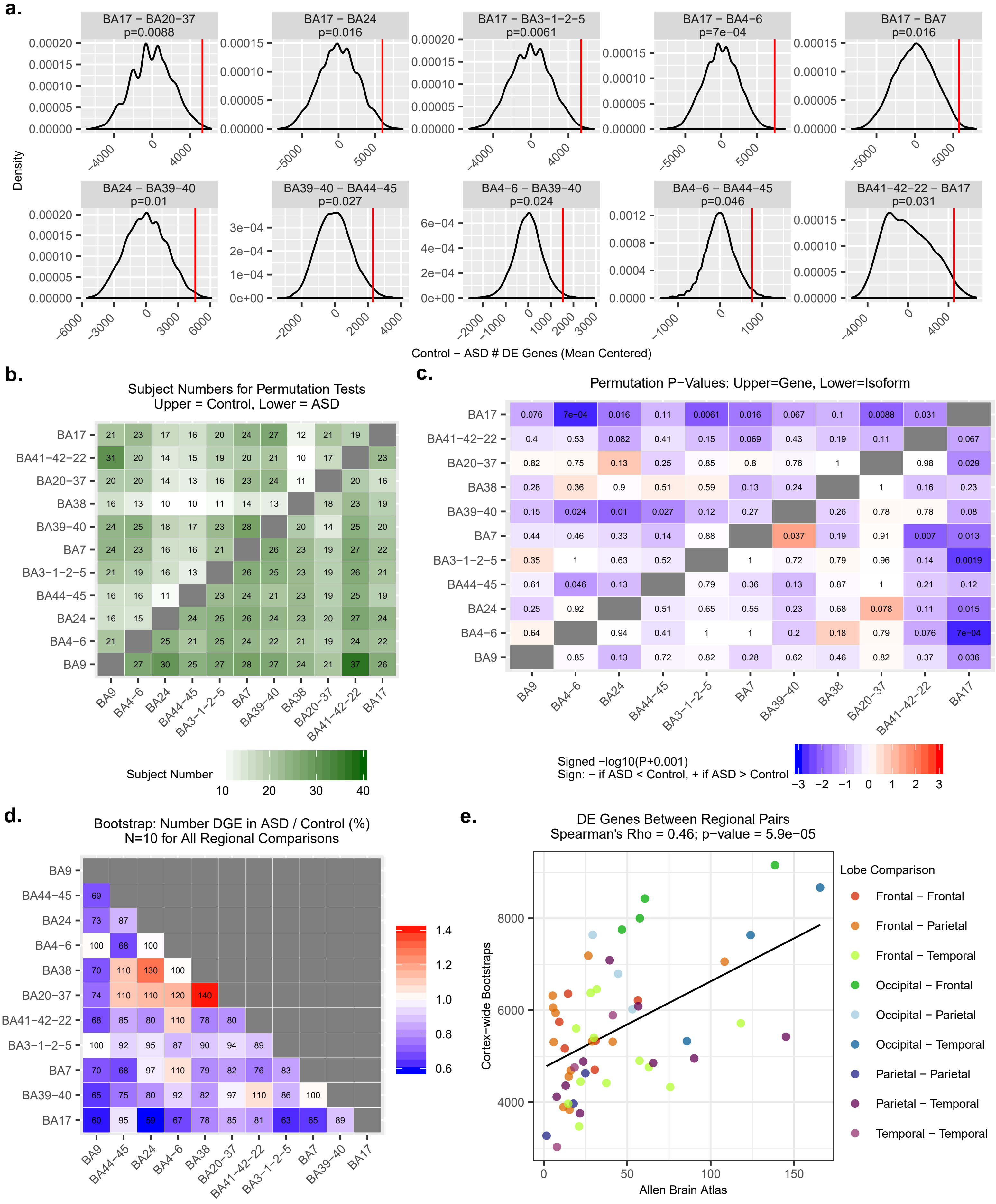


**Extended Data Figure 5 | Transcriptomic Regional Identity Attenuation in ASD. a.** Mean-centered distribution of 10,000 permutations for the significantly attenuated regional comparisons in ASD. Red bar = true difference in the number of DE genes between controls and ASD. **b.** Sample size for all regional comparisons. **c.** Permutation p-values for all regional comparisons. **d.** For 10,000 regional comparison bootstraps, ratio of DE genes in ASD compared to controls. **e.** Number of DE genes between pairs of regions in this study (mean across bootstraps in controls, y-axis) compared to the Allen Brain Atlas (ref. 10, mean across matched regions, x-axis; see Methods for matched regions). This Allen Brain Atlas dataset, with only 2 unique brains, is the best publicly available dataset for comparison (linear least squares regression best fit line plotted).

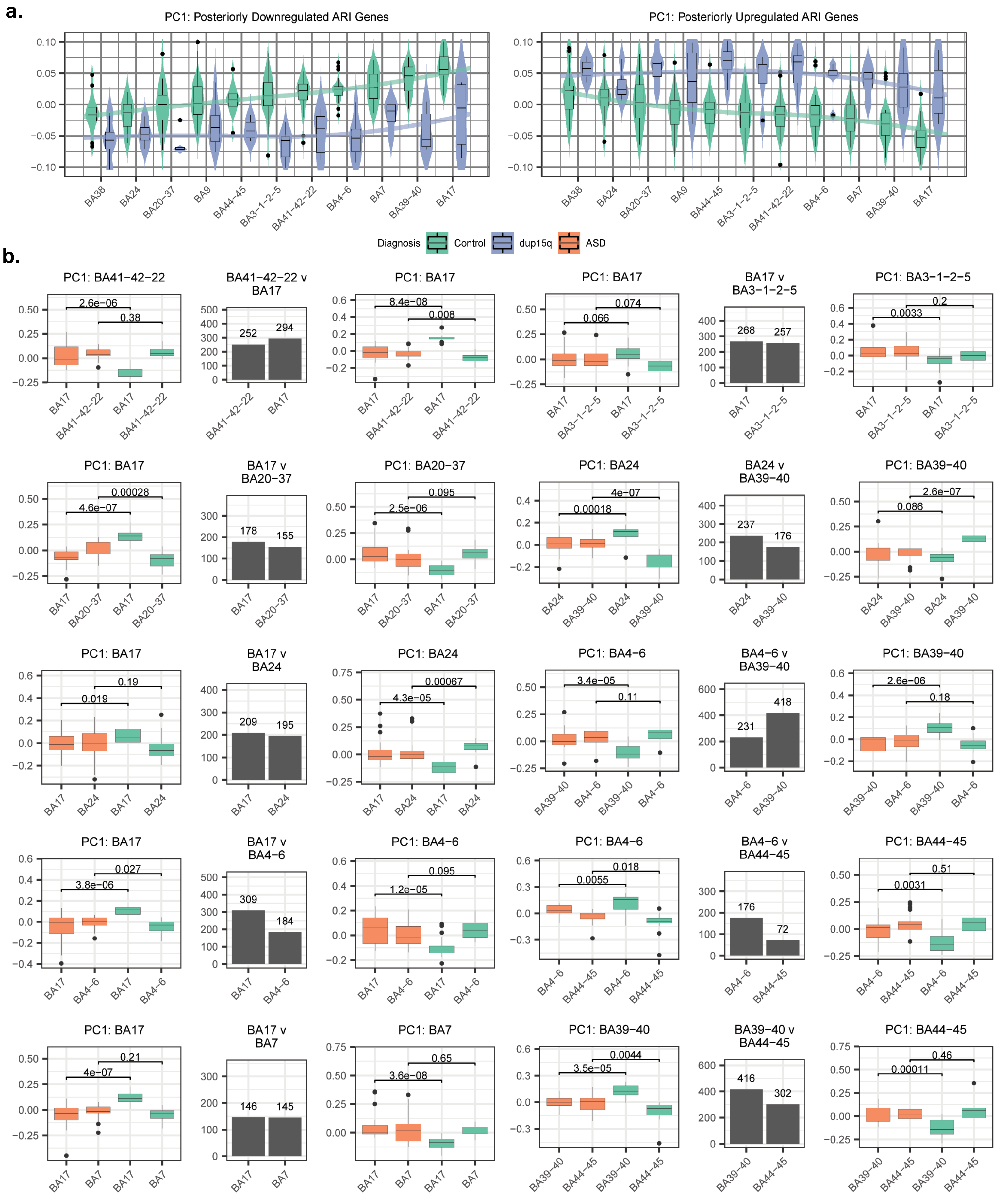


**Extended Data Figure 6 | Additional ARI gene dysregulation. a.** First principal component (PC1) of posteriorly downregulated (1,881, left) and upregulated (1,695, right) ARI genes identified in ASD, plotted in Controls and dup15q (loess regression line plotted). **b.** For each significantly attenuated regional comparison, the identified attenuated regional identity (ARI) genes. At center, number of ARI genes with greater neurotypical expression in each pair of regions. On either side of the barplot, the PC1 of the genes with greater neurotypical anterior (left) or posterior (right) expression is plotted across the pair of regions in Controls and ASD. The Wilcoxon signed-rank test (unpaired) p-value is shown.

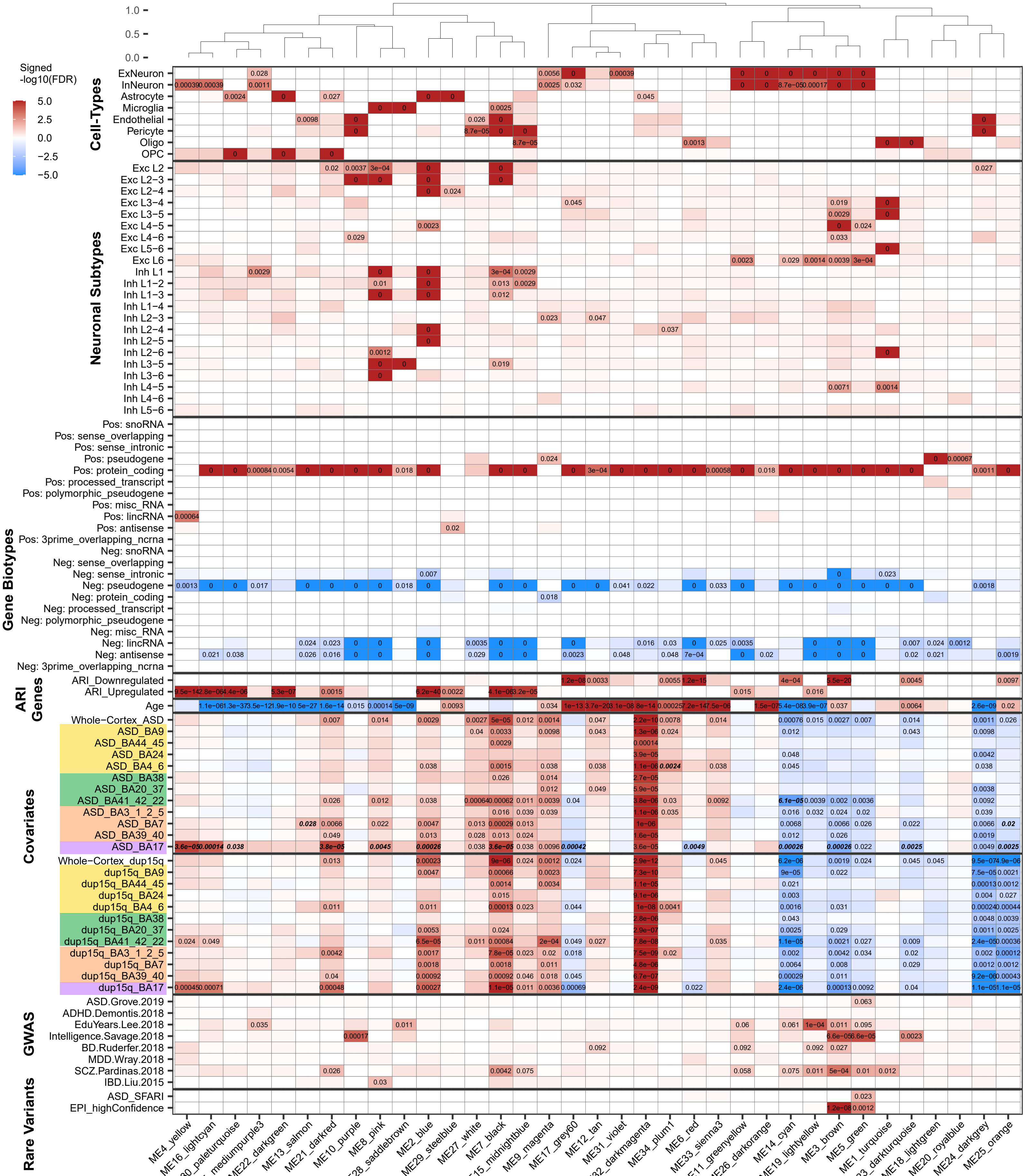


**Extended Data Figure 7 | Gene-level Co-Expression Network Analysis Module Associations.** Top: average-linkage hierarchical clustering of module eigengene biweight midcorrelations. Significant FDR corrected p-values are indicated (FDR < 0.05; for GWAS, FDR < 0.1). Any signed –log10(p) colors greater or less than 5/-5 are set at a max/min of 5/-5 . For ASD, dup15q, and Age covariates, FDR p-value from the linear mixed model testing the association of these covariates with module eigengenes is depicted. For the ASD and dup15q region-specific comparisons, cortical lobule colors are indicated (Fig. 1a), and bold-italic FDR p-values indicate that these regions are effected with significantly greater magnitude than the ASD whole-cortex (Methods). For gene biotypes, both positive and negative enrichment is shown (Methods). Positive enrichment is shown for cell-type, neuronal subtype (ref: Hodge et al, Nature 2019), ARI gene, GWAS, and rare variant enrichment (Methods).

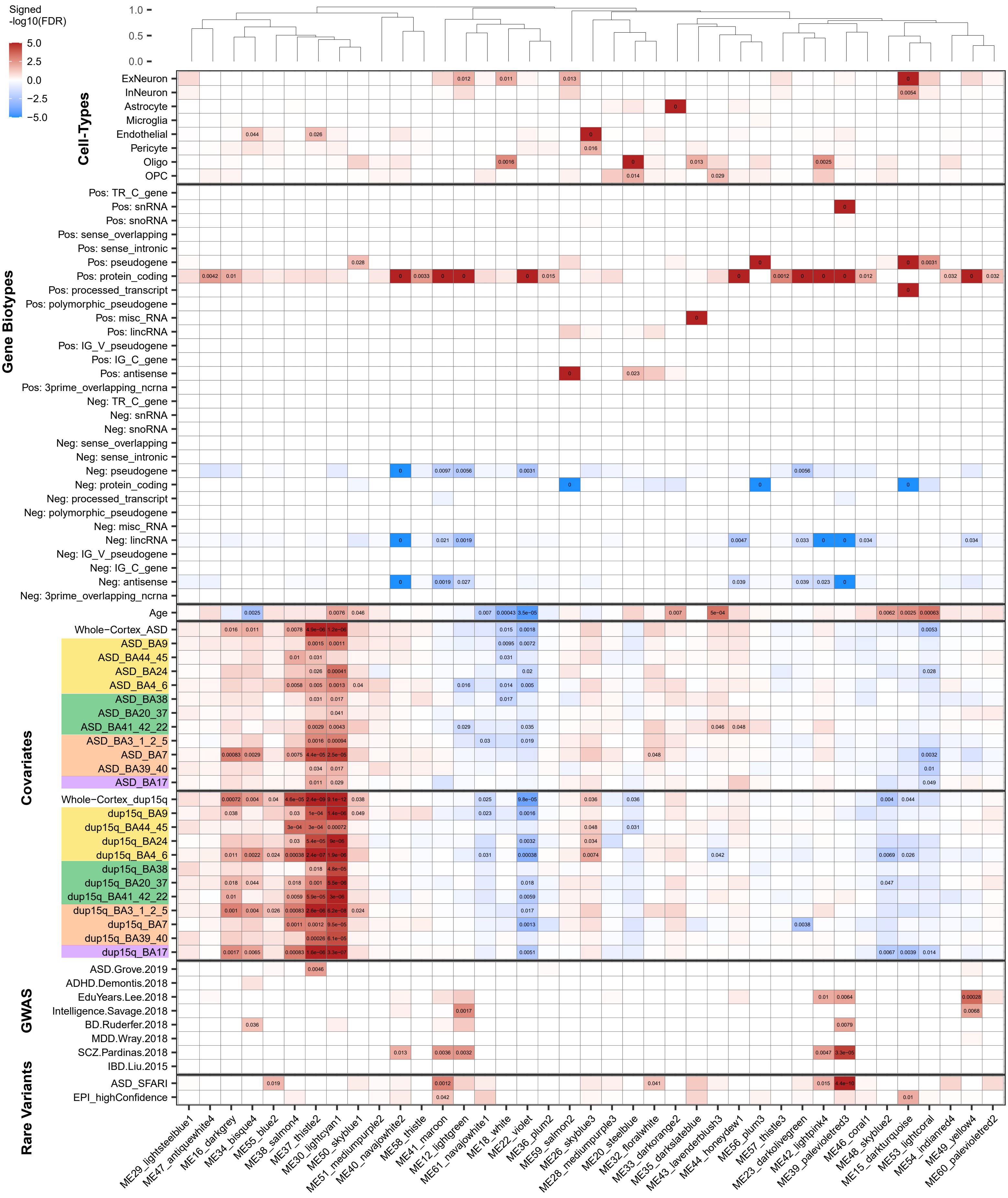


**Extended Data Figure 8 | Isoform-level Co-Expression Network Analysis Module Associations.** Top: average-linkage hierarchical clustering of module eigengene biweight midcorrelations. Significant FDR corrected p-values are indicated (FDR < 0.05; for GWAS, FDR < 0.1). Any signed –log10(p) colors greater or less than 5/-5 are set at a max/min of 5/-5 . For ASD, dup15q, and Age covariates, FDR p-value from the linear mixed model testing the association of these covariates with module eigengenes is depicted. For the ASD and dup15q region-specific comparisons, cortical lobule colors are indicated (Fig. 1a). For gene biotypes, both positive and negative enrichment is shown (Methods). Positive enrichment is shown for cell-type, GWAS, and rare variant enrichments (Methods).

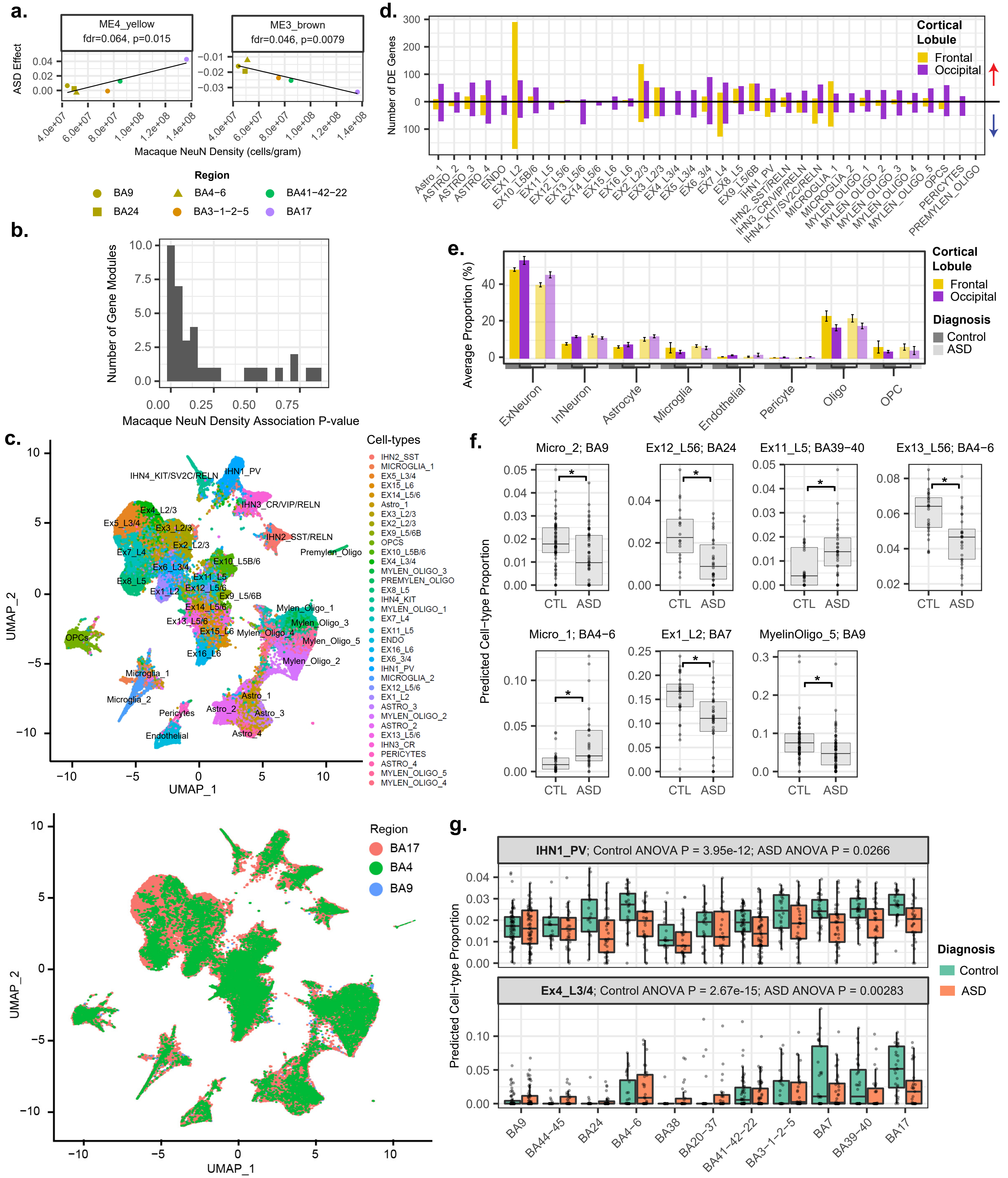


**Extended Data Figure 9 | Neuronal Density Associations, snRNA-seq, and Cell-type Deconvolution. a.** Macaque neuronal density v. module eigengene ASD effect for modules featured in Fig. 4c-d (linear least squares regression). Both p-value and FDR corrected p-value are plotted. **b.** P-value histogram of all gene modules' linear least squares regression with macaque region-specific neuronal density. **c.** UMAP plots of snRNA-seq with cell sub-types (top) and brain regions (bottom) depicted. **d.** Number of genes differentially expressed in ASD in each cell subtype. Upregulated genes are above 0 (red arrow) and downregulated genes are below 0 (blue arrow). **e.** Average proportion of each broad cell-type in each diagnosis x cortical lobule group, derived directly from the snRNA-seq data. **f.** Additional significant (Bonferroni corrected p-value < 0.05) cell-type proportion differences in ASD from cell-type deconvolution. Region and cell-type are indicated in the title of each plot. **g.** For two example cell-types, cell-type proportion attenuation in ASD across regions. ANOVA p-values stratified by diagnosis are shown.

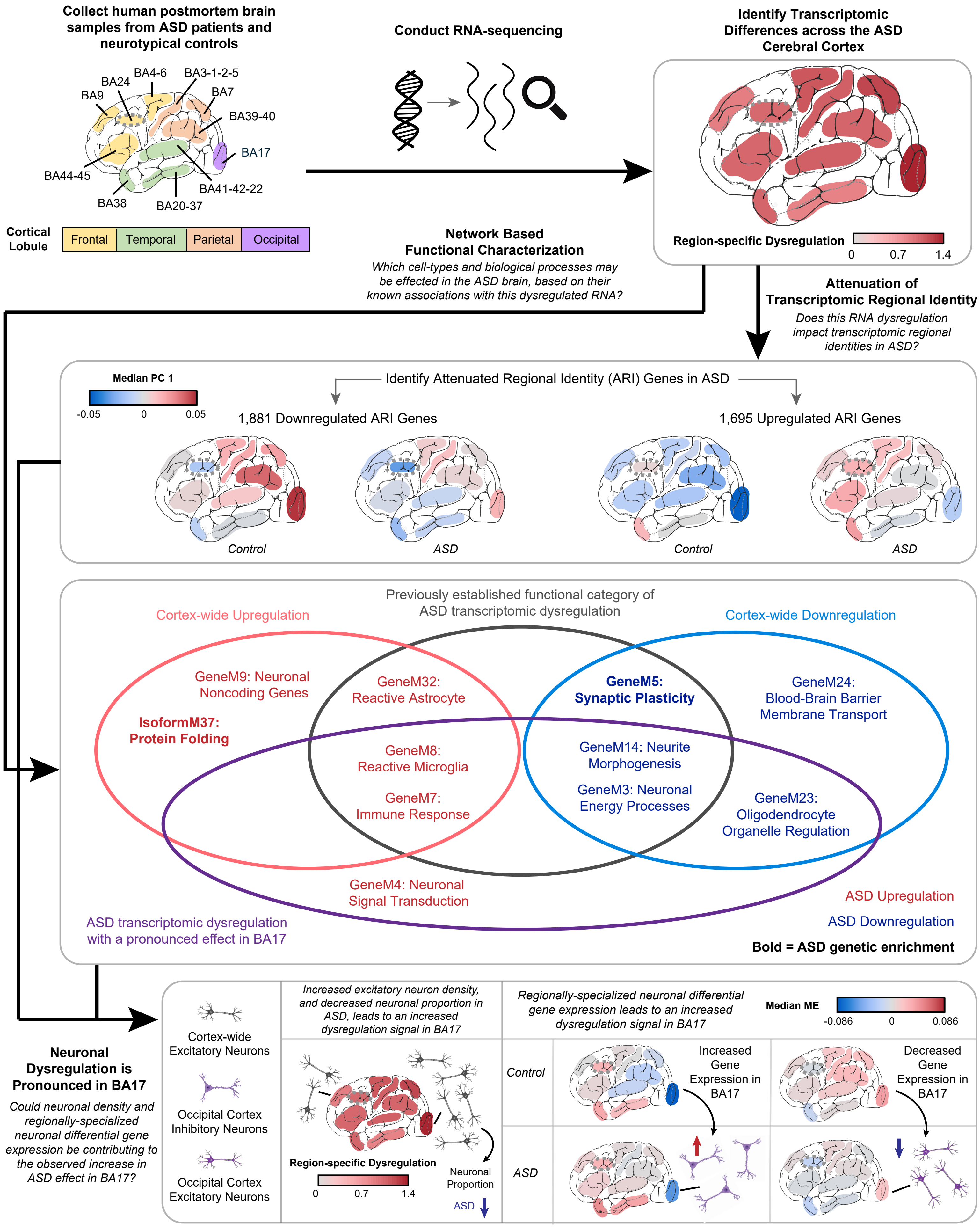


**Extended Data Figure 10 | Results Summary.** Overview of RNA-sequencing experiment and results. Region-specific dysregulation scale in the top right corner and the leftmost portion of the bottom panel depict the region-specific slopes compared to the whole cortex effect from Fig 1d. Median PC 1 of the ARI dysregulated genes is plotted in the middle panel. In the right portion of the bottom panel, the median ME of GeneM4 (left) and GeneM3 (right) is depicted.
