## Supplementary Methods, Supplementary Tables, and Supplementary References for "Broad transcriptomic dysregulation across the cerebral cortex in ASD": SI Guide.docx

**SupplementaryInformation**

*Additional methods (SupplementaryMethods) and detailed descriptions of supplementary table tabs (SupplementaryTables - Descriptions), along with additional references for the supplementary methods (SupplementaryReferences).*

**SupplementaryTable1**

*Metadata and sequencing quality metrics for all samples, along with top gene and isoform expression principal component associations with metadata and sequencing quality metrics.*

**SupplementaryTable2**

*Differential gene expression overlap data, both with previous publications (Voineagu et al. 2011 and Parikshak et al. 2016) and within this dataset (across regions, diagnoses, and whole gene v. isoform datasets). Regional ASD DE gene permutation data is also included.*

**SupplementaryTable3**

*Linear mixed model statistics for biological covariates (diagnosis, region, age, sex), for all genes and isoforms assessed.*

**SupplementaryTable4**

*Statistics from transcriptomic regional identity analysis, including permutation data (for both gene and isoform level expression), bootstrap data, and attenuated regional identity (ARI) gene data. A region matching key for comparing Allen Grain Atlas regions to Brodmann areas is also included.*

**SupplementaryTable5**

*For all genes and isoforms, WGCNA module assignment, kME values, and gene/isoform annotation.*

**SupplementaryTable6**

*Functional characterization data for all gene and isoform modules, along with linear mixed model effects for biological covariates (diagnosis, region, age, and sex) in all modules.*

**SupplementaryTable7**

*Supporting data for the analysis of regionally-variable ASD transcriptomic dysregulation, including: neuronal density and cortical L4 thickness associations with modules, brain area matching keys for these associations, module overlap with low integrity RNA genes, snRNA-seq cell-type proportions across regions (including bootstrapped proportion statistics), snRNA-seq DE gene data, and cell-type deconvolution results and statistics.*
