## Supplementary Methods, Supplementary Tables, and Supplementary References for "Broad transcriptomic dysregulation across the cerebral cortex in ASD": SupplementaryInformation.docx

**Supplementary Information Table of Contents**

Supplementary Methods	1

*Linear model design	1*

*Comparing region-specific ASD effects to whole cortex ASD effects	1*

*ARI gene group formation and functional annotation	2*

*WGCNA network formation and module identification	2*

*Module functional characterization	3*

*Neuronal density and cortical layer 4 association with ASD dysregulation	5*

*snRNA-seq	5*

*Cell-type deconvolution	5*

*Boxplot description	7*

Supplementary Tables (Descriptions)	8

*Supplementary Table 1	8*

*Supplementary Table 2	8*

*Supplementary Table 3	8*

*Supplementary Table 4	9*

*Supplementary Table 5	10*

*Supplementary Table 6	10*

*Supplementary Table 7	10*

Supplementary References	12

### Supplementary Methods

#### **Linear model design**

To select the biological and technical covariates to use in downstream linear mixed-effects models, the EARTH^41^ package in R was used. This package applies the Multivariate Adaptive Regression Splines (MARS) technique to build regression models. The covariates assessed were subject, region, brain bank, diagnosis, sex, age, PMI, sequencing batch, ancestry genotype, and RIN, as well as STAR and Picard Tools RNA-seq quality measures (all listed in **Supplementary Table 1**). For 4 subjects with no recorded PMI, the average of the rest of the subjects’ PMI was used. Before input into the EARTH algorithm, STAR and Picard Tools quality measures were filtered such that collinearity with any other biological or technical covariate was eliminated (only one covariate was kept for every identified collinear pair, with collinearity defined as an adjusted R^2^ > 0.95 between the two covariates). All continuous covariates were centered and scaled for input into the EARTH algorithm and for remaining analyses. A cross-validated approach was used to run EARTH: it was run 10 times with 90% of samples, and then the resulting linear model was tested with the remaining 10% of samples. The median R^2^ across all genes/isoforms was used to assess the performance of each cross-validated EARTH model. Using this metric, the following covariates from the highest performing EARTH model were selected for the gene and isoform linear mixed models used in subsequent transcriptomic analyses:

Gene Model: subject, diagnosis, region, sequencing batch, sex, ancestry, age, age^2^, PMI, RIN, picard_gcbias.AT_DROPOUT, star.deletion_length, picard_rnaseq.PCT_INTERGENIC_BASES, picard_insert.MEDIAN_INSERT_SIZE, picard_alignment.PCT_CHIMERAS, picard_alignment.PCT_PF_READS_ALIGNED, star.multimapped_percent, picard_rnaseq.MEDIAN_5PRIME_BIAS, star.unmapped_other_percent, picard_rnaseq.PCT_USABLE_BASES, picard_alignment.PCT_CHIMERAS^2^, star.uniquely_mapped_percent^2^.

Isoform Model: subject, diagnosis, region, sequencing batch, sex, ancestry, age, age^2^, PMI, RIN, picard_rnaseq.PCT_MRNA_BASES, picard_gcbias.AT_DROPOUT, picard_rnaseq.PCT_UTR_BASES, star.multimapped_toomany_percent, picard_rnaseq.MEDIAN_CV_COVERAGE, picard_insert.MEDIAN_INSERT_SIZE, picard_rnaseq.PCT_INTERGENIC_BASES, picard_rnaseq.PF_BASES.

For both models, ‘subject’ was input as a random effects term (specifically, a random intercept), and diagnosis and region were combined to create one ‘diagnosis x region’ term (eg. ASD_BA17, ASD_BA9, Control_BA17, Control_BA9, etc.). This was done to facilitate region-specific contrasts in downstream analyses. The rest of the covariates were input as fixed effects into the linear mixed models. The ‘variancePartition’^42^ R library was used to visualize the percent of variance explained by each model covariate across all genes/isoforms.

#### **Comparing region-specific ASD effects to whole cortex ASD effects**

To test if region-specific ASD dysregulation was significantly greater in magnitude than the whole cortex dysregulation in the Parikshak et al.^5^ modules, a permutation approach was utilized. The region-specific ASD signed -log_10_(p-value) of each module was tested against a permuted distribution (10,000 permutations) of this statistic generated from randomly assigning cortical regions to samples. Regions were randomly assigned within subjects so that regional sample size was consistent for every permutation and subject variability was controlled. A region was considered significantly more dysregulated than the whole-cortex if the one-tailed p-value derived from comparing the true region-specific ASD signed -log_10_(p-value) to the permuted distribution was less than 0.05. The same approach was implemented to test if the number of region-specific DE ASD genes was significantly greater than the number of whole cortex DE ASD genes, with the number of region-specific DE ASD genes replacing the region-specific ASD signed -log_10_(p-value) as the statistic of interest.

To compare region-specific ASD gene dysregulation effect sizes to the whole cortex ASD effect, we calculated the principal components regression slope comparing the whole cortex ASD log_2_ Fold Change (FC/effect) to the region-specific ASD log_2_ FC for the 4,223 genes identified as DE in ASD across the whole cortex. We then generated a bootstrapped distribution (1,000 bootstraps) for each of the 11 region-specific slopes (sampling with replacement from the region of interest for each ‘diagnosis x region’ group) to calculate a 95% confidence interval for these slopes. Sample size was kept consistent for each bootstrap with the number of samples from each ‘diagnosis x region’ group.

#### **ARI gene group formation and functional annotation**

To evaluate the ARI genes across the whole-cortex, instead of only in the regional pairs in which they were identified, the ARI genes from regional pairs containing either BA17 or BA39-40 were assembled into two groups: the union (without duplicates) of ARI genes with higher Control expression in BA39-40 and BA17 relative to other regions (posteriorly ASD-downregulated ARI genes), or the union (without duplicates) of ARI genes with higher Control expression in the remaining cortical regions relative to BA39-40 and BA17 (posteriorly ASD-upregulated ARI genes). Genes which were sorted into both groups (eg. highest expression in BA39-40 v. BA44-45 in one regional comparison, and highest expression in BA7 v. BA17 in another) were removed. Additionally, for each remaining ARI gene, the median Control gene expression in BA17 and BA39-40 (from the regressed gene expression dataset used for the permutation analysis, using all Control samples) was compared to the median across all remaining regions. Only ARI genes with higher median expression in their respective group (eg. higher median expression in BA17 and BA39-40 in the posteriorly ASD-downregulated ARI gene group) were retained. For each gene in each of the two groups, the linear contrast comparing BA17 and BA39-40 gene expression to all other cortical regions was assessed in Controls with the same linear model workflow and normalized, outlier-removed gene expression dataset used to identify DE genes and isoforms described before. The beta values and p-values from this analysis are shared in **Supplementary Table 4** and for top attenuated transcription factors (TFs) in **Figure 2c-d**.

To functionally characterize the ARI gene groups, we performed cell-type and gene ontology enrichment, identified transcription factors present, and calculated transcription factor binding site enrichment. Cell-type enrichment was conducted with EWCE,^43^ with broad (Level 1) neural cell-type gene markers acquired from Lake et al. Nat Biotechnol 2018^4^^4^ (frontal and visual cortex samples combined). To obtain cell-type specificity scores, first genes were filtered such that the gene needed to have a mean UMI of 0.005 across all cells. Then, gene UMI averages were taken across all Level 2 cell-types, and these averages were used to generate the cell-type specificity scores utilized by EWCE to calculate cell-type enrichment in the ARI gene groups. This approach was taken to reduce bias introduced by differing numbers of cells across Level 2 cell-types when calculating Level 1 cell-type specificity scores. 100,000 bootstraps were generated to determine cell-type enrichment with EWCE. gProfileR^4^^5^ was used for gene ontology enrichment, with FDR-adjustment for p-values, strong hierarchical filtering, and a required overlap size of 10 genes. For the ARI downregulated gene group, a max set size of 2500 was enforced, whereas no max set size was enforced for the ARI upregulated gene group. Only ‘BP’ (biological process) terms were included in **Figure 2** and **Supplementary Table 4**. Transcription factor binding site enrichment was also conducted with gProfileR,^4^^3^ with a Bonferroni-adjustment for p-values and strong hierarchical filtering. To identify transcription factors within the ARI gene groups, AmiGo 2^4^^4^ was used to acquire all genes in GO:0003700 (DNA-binding transcription factor activity) in the Homo sapiens organism (Gene Ontology Consortium,^47,4^^8^ accessed May 7, 2020).

#### **WGCNA network formation and module identification**

Weighted Gene Co-Expression Network Analysis (WGCNA)^10^ was conducted to sort observed gene and isoform expression dysregulation into empirically-informed networks which could provide precise functional insight into affected neural cell-types and biological processes. Regressed gene and isoform expression datasets containing only the random effect of subject, the fixed biological effects (diagnosis, region, age, age^2^, sex, and ancestry), and the model residual were used for WGCNA signed network generation. Regression was performed as described previously for the previously identified co-expression modules. A soft-threshold power of 6 was chosen for gene network generation, whereas a power of 10 was selected for isoform network generation. These values were selected to optimize induced scale-free topology in the gene and isoform networks (R^2^ > 0.8). For the gene-level WGCNA, a robust version of WGCNA (rWGCNA)^21^ was implemented to mitigate the influence of potential sample outliers in network formation. Subjects within each diagnosis group were randomly selected (with replacement) for inclusion in the adjacency matrix (formulated using the bi-midweight correlation of genes) and subsequent TOM matrix generation, 100 times. These TOMs were merged into one consensus TOM through first using a quantile scale of 0.95 to calibrate each TOM, and then taking the median across all TOMs to create the consensus TOM. To identify modules from the consensus TOM, the ‘cutTreeHybrid’ function was used with average linkage hierarchical clustering of the consensus TOM, a deep split of 4, cut height of 0.9999, a negative PAMstage, and minimum module size of 50. Modules within a cut height of 0.1 were merged.

Since rWGCNA could not be implemented for the isoform expression data due to memory allocation limitations, the ‘blockwiseModules’ function was used with 4 blocks (26,000 or less isoforms per block) to generate the isoform network and identify modules. The same module identification parameters (except for the soft power threshold) used for the gene network were also used for the isoform network. To test the robustness of the isoform network, a permutation approach was utilized.^22,4^^9^ For each module, this method tests if the mean connectivity within the module (also defined as the module’s density, or the average intramodular topological overlap) is significantly different from that of modules of equivalent size randomly selected from the same network (n=5,000 permutations). One-tailed p-values were calculated through comparing the permuted distribution to the true mean connectivity for each module, and only modules with p-values < 0.05 were retained. When merging modules from all blocks for the isoform network, a merge cut height of 0.2 was used.

Module eigengenes (MEs) were calculated for all modules using the regressed gene and isoform expression dataset used to generate the networks. We only retained isoform modules which were non-redundant with gene modules forward for further analysis. To achieve this, isoform and gene MEs were clustered using the ‘cutTreeHybrid’ WGCNA^10^ function using average linkage hierarchical clustering of the bi-midweight correlation of the MEs, a deep split of 4, a negative PAMstage, a minimum module size of 1, and a cut height of 0.9999. Any isoform modules which clustered with gene modules were labeled as overlapping with the gene modules, with the exception of Isoform_M26_skyblue3, which upon visual inspection was suitably distant from the other gene modules within its cluster to be considered distinct. To determine if any of these other overlapping isoform modules were distinct enough from the gene modules to be retained for further analysis, for each of the conserved isoform modules an over-representation analysis (ORA) was conducted with each of the gene modules in its identified cluster. Any isoform modules which had no significant overlap (p > 0.01) were retained for further analysis, of which only two were identified - Isoform_M55_blue2 and Isoform_M61_navajowhite1. In total, 39 distinct isoform modules were carried forward for further analysis out of the original 61 identified isoform modules.

#### Module functional characterization

Gene and the distinct isoform MEs were assessed with the gene and isoform linear mixed models containing all of the biological covariates from the full models described previously (the technical covariates were not included, since these covariates were previously removed from the regressed expression data used to generate the MEs). The same limma^3^^5^ workflow was implemented as described before for calculating DE genes and isoforms. Whole cortex and region-specific ASD and dup15q effects were also ascertained as described previously for the DE gene and isoform analysis. A covariate required an FDR-adjusted p-value < 0.05 to be considered associated with any ME. To determine if any region-specific ASD effects in the gene MEs were significantly greater than the whole cortex ASD effect, a permutation approach was used which was synonymous to the previously described method used with the Parikshak et al. MEs. Regionally-variable modules are those with any region having a region-specific ASD effect significantly greater than the whole cortex ASD effect (p < 0.05).

To further functionally characterize modules, we calculated enrichments for neural cell-types, neuronal subtypes, gene ontology terms, protein-protein interactions, the ARI gene groups, gene biotypes, relevant GWAS, ASD and epilepsy associated rare variants, and gene modules previously associated with ASD published in Parikshak et al. Nature 2016^5^ and Gandal et al. Science 2018b.^1^ While all of these enrichment analyses were performed for the gene modules, only a subset were performed for the isoform modules (neural cell-types, gene ontology terms, gene biotypes, psychiatric GWAS, and ASD and epilepsy associated rare variants).

Neural cell-type enrichment was performed with EWCE^43^ as previously described for the ARI gene groups. For neuronal subtype enrichment, medial temporal gyrus single neuron RNA-seq from the Allen Brain Map^13,^^43^ was used to define neuronal subtype specific markers for enrichment analysis with EWCE.^43^ EWCE was implemented as previously described for the ARI gene groups, with the Allen Brain Map neuronal cells being grouped into cortical layer groups (eg. Exc L2, Inh L2-3), for cell-type enrichment. For gene ontology terms, the Metascape^50^ web portal was used with default functions (‘Express Analysis’). Only ‘GO Biological Process’ terms with an FDR-adjusted p-value < 0.05 were examined for each module. PPI annotations and enrichments were calculated with STRING,^51^ run with default settings in June 2019. A direct connection FDR-corrected p-value < 0.05 was needed for a module to be considered significantly enriched with PPIs. ARI gene group enrichment was calculated with ORA, with an FDR-corrected p-value < 0.05 and OR > 1 being required for a significant enrichment.

Gene biotype enrichment was determined with a permutation approach. The number of each unique gene biotype was first acquired for each module. Then, for each permutation (10,000 in total) gene biotypes were sampled across all genes without replacement and randomly assigned. The number of each unique gene biotype in each module was collected for each permutation. A distribution could then be created for each unique gene biotype in each module across the 10,000 permutations. Both over- and under-enrichment of each unique gene biotype in each module was determined directly with this distribution (one-tailed p-value). An FDR-corrected p-value < 0.05 was required for a significant enrichment.

For the psychiatric GWAS enrichments, partitioned heritability was calculated with stratified LD Score regression^52^ (run with recommended settings) using 10 kb windows around genes (matched genes were used for isoform modules). An FDR-corrected p-value < 0.1 was required for a significant GWAS enrichment (the threshold for significance was relaxed since many of the best available GWAS datasets utilized are underpowered, particularly the ASD GWAS). We selected the most recent and best powered GWAS which were relevant and interesting for comparison with these gene and isoform modules, including GWAS conducted for ASD,^11^ ADHD,^53^ BD,^5^^4^ MDD,^5^^5^ SCZ,^5^^6^ Educational Attainment,^5^^7^ Intelligence,^5^^8^ and IBD.^59^ Logistic regression was used for rare variant enrichment, controlling for both gene length and GC content, with an FDR-corrected p-value < 0.05 being required for a significant enrichment. Syndromic and highly ranked (1 and 2) ASD SFARI^12^ gene and high-confidence Epilepsy (compiled by D. Polioudakis et al. Neuron 2019)^60^ gene associations were examined. Finally, ORA was used to assess previous module enrichment, with an FDR-corrected p-value < 0.05 and OR >1 indicating a significant positive overlap.

#### **Neuronal density and cortical layer 4 association with ASD dysregulation**

A linear model was used to compare region-specific macaque NeuN density^15^ to region-specific ASD effects (model beta) in the regionally-variable gene MEs. Macaque brain areas were matched to Brodmann areas (shared in **Supplementary Table 7**), with six regions matching between this dataset and the macaque dataset. FDR-corrected p-values < 0.1 were considered significant neuronal density associations (the FDR threshold was relaxed, since only 6 regions/points were available for every comparison). A leave-one-out cross-validation was performed to assess individual regional contributions to neuronal density associations, in which a single region was withheld and linear model statistics were re-calculated. In addition to neuronal density, we also examined the association between cortical layer 4 thickness^18^ (von Economo and BigBrain estimates, as shared in the publication) and region-specific ASD effects in the regionally-variable gene MEs. All 11 regions were matched to layer 4 thickness measures (this key is shared in **Supplementary Table 7**). This comparison was also performed with a linear model, with FDR-corrected p-values < 0.05 considered significant layer 4 thickness associations.

#### **snRNA-seq**

Control and ASD samples (8 total) matched for co-variates (i.e age, sex, manner of death) were processed in the same nuclear isolation batch to minimize potential batch effects. These subjects included: UMB5144 BA17 and BA9, AN08792 BA17 and BA4-6, AN10679 BA17 and BA4-6, UMB4787 BA17 and BA4-6. 50 mg of sectioned brain tissue was homogenized in 2.5 mL of RNAase-free homogenization buffer (250mM sucrose, 5mM MgCl2, 25mM KCL, 10mM Tris pH8, 1 uM DTT, 0.2U RNaseIN, 1% BSA, 0.01% Triton X-100, 0.001% Digitonin in RNAse-free water) using glass dounce homogenizer on ice. The homogenate was filtered and subjected to a two layer micro-iodixanol nuclei centrifugal gradient (50%/30%) for 13500g for 20 minutes at 4°C. Supernatant was carefully removed and the nuclei containing pellet were resuspended in RNase-free PBS pH7.4, 5mM MgCl2, 1% BSA, 0.2U RNaseIN. The nuclear suspension was filtered twice through a 30 um cell strainer. Nuclei were counted using a hemocytometer and diluted to 1,000 nuclei/uL before performing single-nucleus isolation on the 10X Genomics controller.; The 10X capture and library preparation protocol was used without modification. Single-nucleus libraries from individual samples were pooled and sequenced on the NovaSeq 6000 machine (average depth 60,000 reads/nucleus).

Raw snRNA-seq data processing was performed with 10X Genomics CellRanger software, Seurat,^61^ and Liger.^62^ CellRanger was used with default parameters, except we utilized the human pre-mRNA reference file (ENSEMBL GRCh38)^27^ to insure capturing intronic reads originating from pre-mRNA transcripts abundant in the nuclear fraction. Individual libraries were analyzed in Seurat for quality control metrics and filtering. Individual libraries were filtered to retain nuclei with at least 500 genes expressed and less than 5% of total UMIs originating from mitochondrial RNAs. Individual matrices were combined, UMIs were normalized to the total UMIs per nucleus and log transformed. Nuclei for all ASD and control subjects from both the PFC and OCC were used for clustering with integrative non-negative matrix factorization with K=40 and lamba= 5.0 followed by quantile normalization and louvain clustering in Liger. We then visualize integrated cells in two-dimensional space with Uniform Manifold Approximation and Projection (UMAP).

#### **Cell-type deconvolution**

**Selected datasets**

Bulk RNA-seq: For the 808 samples from 11 regions, we used the residuals of the regression of the bulk data against technical covariates. While we ran the deconvolutions for all samples, we explicitly excluded the Dup15q samples from the downstream comparisons between the ASD and CTL cohorts.

Single-nuclei RNA-seq: The single-nuclei dataset used in this analysis was obtained from the frontal cortex (FC) and primary visual cortex (V1C) of 4 individuals (2 ASD and 2 CTL) overlapping with the bulk tissue cohort, comprising four FC libraries and four V1C libraries (described in the preceding section). Cell type assignments for the bulk-tissue-overlapping snRNA-seq libraries were obtained by looking at the expression of canonical markers from years of culminated mouse and human studies and recent single-cell atlases. Specifically, we utilized the Hodge 2019 (Allen Institute)^13^ and Lake 2018^18^ papers to establish frontal cortex and V1 specific signatures found previously. Based on both the expression level of the gene, but also the percentage of cells within a cluster that expressed said gene, we identified 35 cell types/states. BA17 specific neurons are superficial neurons in V1 that express SYT2 and RORB higher than frontal cortex, and Layer 4 V1 Ext neurons express PHACTR2 and EYA4 over other areas.

Here are examples of core genes used to establish excitatory neuronal types: SLC17A7, RBFOX3. For Excit L2/3: “LAMP5",“CUX2”,“GLRA3", “CUX1”, “LHX2",“CBLN2”, “RASGRF2", “COL5A2”, “LMO3", “SATB2”. For Excit 3/4: “RORB”, “PCP4”, “LMO3", “CUX2”, “SATB2", “NEFM”, “PHACTR2",“EYA4” . For Excit L5/5B: “BCL11A”,“CRYM”,“FOXP2",“BCL11B”,“FEZF2", “RORB”, “DKK3", “TLE4”, “SEMA3E”, “LMO4", “CCK”,“ETV1", “NEFH”, “CNTN6", “FOXO1”, “OPN3", “LIX1”, “SYT9", “S100A10”, “LDB2", “CRIM1”, “PCP4", “SATB2”, “CRYM”. For Excit L6: “GLRA3", “LMO3”, “BHLHE22", “RORB”, “NNAT”, “FOXP2”, “ETV1", “FEZF2”, “TLE4", “GRIK4”, “NTNG2", “OPRK1", “NR4A2”, “BCL11B”, “THY1”.

**Data processing for deconvolution analyses**

We applied the following processing steps to the bulk tissue RNA-seq and snRNA-seq data prior to running CIBERSORTx.

1. **Bulk RNA-seq:** For the post-regression residual matrices, gene names were converted from ENSEMBL IDs to HGNC symbols. Any genes that were not mapped (by *biomaRt* in R) were removed. Gene expression values were transformed from their log-base-2 values to non-log expression values, as required by CIBERSORTx.
2. **snRNA-seq:**
   a. Expression analysis: The input data format consisted of standard *CellRanger* output matrices. The matrices were read into *Seurat* (https://cran.r-project.org/web/packages/Seurat) objects in R. Lenient QC cutoffs of “Number of RNA features > 500” and “Percent Mitochondrial genes < 10” were chosen to filter out cells. The *Liger* cell-type-cluster labels were associated with cells, and any cells without identified cell types were removed from the analysis. We again used *scran* but modified the previous pipeline slightly: (I) removed cell types that have < 10 cells within the sample; (II) found the size factors using pool sizes of 20, 30, 40, 50, and 60; and (III) further removed any cells that produced negative size factors. Finally, the counts for each cell were converted to a weighted counts-per-million scale as $Weighted\_CPM\left( cell i, gene g \right)= {10}^{6}.Size\_factor\left( cell i \right).\frac{Counts\left( cell i, gene g \right)}{\sum_{g=1}^{N_{genes}} Counts\left( cell i, gene g \right)}$, and $N_{genes}$ = Total number of genes.
   b. Pooling of cells: We combined cells from the FC and V1C regions into a single reference matrix for the deconvolution. Each of the 4 FC and 4 V1C libraries was run through the preprocessing steps separately and combined subsequently. The merging of matrices was carried out by concatenating *pandas* data frames in Python, using a join on overlapping gene names. It is worth noting that this merging process results in only the intersecting genes across all datasets being included, and due to the removal of rows corresponding to non-overlapping genes, the resulting cell expression vectors may not be normalized to ${10}^{6}$. The numbers of cells included in the final deconvolution analysis are 145,373 cells.

**CIBERSORTx parameters**

We downloaded the *fractions* module of CIBERSORTx (doi.org/10.1038/s41587-019-0114-2) from the website, in the form of a Singularity image of the Docker file. We ran the program in the mode that accepts a full single cell count matrix as input, thereby implicitly generating a reference matrix (the parameter *single_cell* is set to *TRUE*), and set the batch-correction mode to the ‘S’ mode (the parameter *rmbatchSmode* is set to *TRUE*).

**Testing for differences in cell fractions between ASD and CTL groups**

To evaluate whether differences, between the ASD and CTL groups, in the cell fractions of particular cell types in each region were statistically significant we calculated p-values from the Wilcoxon Rank-Sum Test and two-sided Kolmgorov-Smirnov (KS) Test on the distributions of the two groups. These were calculated using the *scipy.stats.ranksums* and *scipy.stats.ks_2samp*, respectively in python’s *scipy* library. For multiple-hypothesis-testing correction, we performed Bonferroni correction for each region: that is, we divided the p-values for each cell type and each region by the number of cell types considered (and not by the product of the number of cell types and the number of regions). Wilcoxon rank-sum test p-values that were lower than the Bonferroni corrected significance threshold (0.05/35 = 0.0014) were considered to represent significant differences in cell-type proportion.

**Testing for differences in cell fractions across regions**

ANOVA was used to test for differences in cell fractions across all eleven regions separately in ASD and CTL groups. Significant differences were those lower than the Bonferroni significance threshold, corrected across all cell-types within diagnosis groups (0.05/35 = 0.0014). ASD differences were considered attenuated if the ASD p-value was greater than that of the CTL group.

#### Boxplot Description

For all boxplots in this manuscript, the median is the center line, box limits are the upper and lower quartiles, whiskers extend to the 1.5x interquartile range, and points beyond the whiskers are outlier observations.

### Supplementary Tables

#### **Supplementary Table 1**

*datMeta_datSeq*

Available metadata and sequencing quality metrics for all samples from the processed dataset (outliers which were removed are not included).

*R2_exprPCs_Covariates*

Data for Extended Data Figure 2c. Top 15 principal components (PCs) for processed (normalized and outliers removed) gene and isoform expression data. Percent expression variance explained by each PC is indicated. Cells are adjusted R^2^ values from the linear model `PC ~ Covariate`.

#### **Supplementary Table 2**

*Voineagu_Parikshak_byRegion*

Statistics for the region-specific ASD covariate in previously discovered ASD co-expression modules examined in Main Figure 1b. Permutation p-values, to assess if the region-specific ASD effect is significantly greater than the whole cortex ASD effect (Methods), are also calculated for the Parikshak et al. 2016^5^ modules with an FDR < 0.05.

*ASD_DEGene_Regional_Permutation*

Number of DE genes (FDR < 0.05) for each region-specific ASD contrast, along with Permutation p-value (to assess if the region-specific number of ASD DE genes is significantly greater than the whole cortex number of ASD DE genes; Methods).

*ASD_DE_Regional_Overlap*

Overlap in region-specific ASD DE genes and isoforms (FDR < 0.05) between regions (where ‘whole cortex’ can be considered its own region). For whole cortex comparisons, the principal components slope with 95% CI (obtained from a bootstrapped distribution; Methods) from Main Figure 1d is also listed.

*ASD_DE_GeneIsoform_Overlap*

For each set of region-specific ASD DE genes and isoforms (FDR < 0.05), overlap between DE genes and DE isoforms.

*ASD_dup15q_DE_Overlap*

Overlap in ASD DE genes/isoforms and dup15q DE genes/isoforms (FDR < 0.05) for each region-specific comparison.

#### **Supplementary Table 3**

*DEGene_Statistics*

For all genes assessed, gene annotation and DE statistics for biological covariates.

*DEIsoform_Statistics*

For all isoforms assessed, gene annotation and DE statistics for biological covariates.

#### **Supplementary Table 4**

*TRI_Gene_Permutation*

Statistics for the gene-level permutation analysis to determine if transcriptomic regional identity is altered in ASD. Regional comparison, permutation p-value, and sample size (N) for ASD and control subjects is indicated. ‘CTL_NumDGE - ASD_NumDGE` is the difference in the number of DE genes between regions for control and ASD subjects (a positive number indicates more DE genes between regions in controls, and a negative number indicates more DE genes between regions in ASD). This difference is adjusted with the mean of the permuted distribution (true difference – permutation mean).

*TRI_Isoform_Permutation*

Statistics for the isoform-level permutation analysis to determine if transcriptomic regional identity is altered in ASD. Regional comparison, permutation p-value, and sample size (N) for ASD and control subjects is indicated. ‘CTL_NumDGE - ASD_NumDGE` is the difference in the number of DE genes between regions for control and ASD subjects (a positive number indicates more DE genes between regions in controls, and a negative number indicates more DE genes between regions in ASD). This difference is adjusted with the mean of the permuted distribution (true difference – permutation mean).

*TRI_Bootstrap*

Statistics for bootstrap analysis to gain a general sense of transcriptomic regional identity in control and ASD subjects. Mean DE genes between regions, standard deviation, and 95% confidence interval for the bootstrapped distribution for each regional comparison is listed separately for ASD and control subjects.

*AllenBrainAtlas_RegMatch*

Key for comparing Allen Brain Atlas^9^ regions to Brodmann Areas, which was used for Extended Data Figure 5e.

*All_ARI_Genes*

For each attenuated regional comparison, identified attenuated regional identity (ARI) genes are listed with gene annotation information. ‘Wilcoxon_PValue’ is the paired Wilcoxon signed-rank test p-value for the difference in gene expression between regions for the control subjects (with these controls being the same controls used for the TRI permutation analysis). ‘ARI_Gene_Group’ status is also indicated, if this gene passed filters to be included in the downregulated (BA17_BA39-40; Main Figure 2c) or upregulated (Other_Cortical_Regions; Main Figure 2d) ARI gene group. ‘TF_Status’ is ‘yes’ if the gene is a known transcription factor, and ‘no’ otherwise. The FDR-corrected p-value and logFC from the contrast of the `BA17_BA39-40` group v. ‘Other_Cortical_Regions’ group in all of the Control subjects is also included, labeled as ‘Group_Contrast_FDR’ and ‘Group_Contrast_logFC’, respectively.

*ARI_Gene_Group_Enrichment*

Broad neural cell-type and any significant transcription factor binding site enrichments for ARI gene groups. FDR-corrected p-values are indicated.

*ARI_Gene_Group_GO*

For each ARI gene group, up to 10 gene ontology terms enriched in the module (ranked by the FDR corrected p-value) are shown. ‘InModule_InGOTerm’ shares how many genes in the module are in the GO term (left), with the number of genes in the GO term on the right.

#### **Supplementary Table 5**

*Gene_Level*

WGCNA module assignment and kME values, along with gene annotation, for all genes assessed.

*Isoform_Level*

WGCNA module assignment and kME values, along with gene annotation, for all isoforms assessed.

#### **Supplementary Table 6**

*GeneModules*

Functional characterization of all gene modules. For neural cell-type, neuronal subtype, gene biotype, ARI gene, GWAS, rare variant, and psychiatric co-expression module enrichments, FDR corrected p-values are displayed. For linear model covariates utilized to assess biological covariate effects in module eigengenes, beta (linear model effect) and FDR corrected p-values are both shared.

*IsoformModules*

Functional characterization of all gene modules. For neural cell-type, gene biotype, GWAS, and rare variant enrichments, FDR corrected p-values are displayed. For linear model covariates utilized to assess biological covariate effects in module eigengenes, beta (linear model effect) and FDR corrected p-values are both shared.

*GeneModule_Ontology*

For each gene module, up to 10 gene ontology terms enriched in the module (ranked by the FDR corrected p-value) are shown. ‘InModule_InGOTerm’ shares how many genes in the module are in the GO term (left), with the number of genes in the GO term on the right.

*IsoformModule_Ontology*

For each isoform module, up to 10 gene ontology terms enriched in the module (ranked by the FDR corrected p-value) are shown. ‘InModule_InGOTerm’ shares how many isoforms in the module are in the GO term (left), with the number of genes in the GO term on the right.

#### **Supplementary Table 7**

*NeuronalComposition*

Macaque NeuN regional density (Collins et al. 2010^15^) and L4 cortical thickness (Wagstyl et al. 2020^18^) association with the ASD effect of gene module eigengenes, and regionally-specific neuronal cell-type (Lake et al. 2018^18,4^^1^) enrichment in gene modules. Leave-one-out cross validation results are included for macaque NeuN associations with module eigengenes (region left out is indicated in column names). Cell-type enrichment cells display the FDR corrected p-value.

*Macaque_BA_Match*

Key for comparing the macaque brain areas from Collins et al. 2010^15^ with Brodmann Areas, used for the macaque NeuN regional density comparisons with ASD module eigengene effects.

*vonEconomo_BA_Match*

Key for comparing von Economo brain areas with Brodmann areas, used for the L4 thickness comparisons with ASD module eigengene effects.

*LowGIS_Enrichment*

GSEA^6^^3^ was used to determine if known low integrity genes with increased susceptibility to postmortem degradation^6^^4^ were enriched in any gene or isoform modules. Specifically, the top 1,000 genes with the lowest GIS scores as established in Feng et al.^6^^4^ were used. For GSEA, kME values were used to order module genes (highest kME to lowest kME). None of the enrichments were significant (FDR corrected p-value less than 0.05). FDR and Bonferroni multiple comparisons correction was conducted separately with gene and isoform modules.

*snRNAseq*

Both broad (top) and sub- cell-type (bottom) average proportions (given as percents) and numbers of up- and down-regulated genes are listed, stratified by cortical lobule (frontal = BA9, BA4_6; visual = BA17) and diagnosis (Control and idiopathic ASD). Standard errors (SE) are also included for cell-type proportions. Average cell-type proportions and standard errors were scaled such that each Lobule x Diagnosis group sums to 100.

*snRNAseq_scDC*

Results from scDC^38^ bootstrapping of cell-type proportions. For the statistics above the black line, astrocytes were the reference cell-type and control, frontal cortex samples were the reference diagnosis x region group. For the statistics below the black line, astrocytes were the reference cell-type and control, occipital cortex samples were the reference diagnosis x region group. Interaction effects should be interpreted with respect to these reference groups.

*cellTypeProportions*

Cell-type proportions for all bulk RNA-seq samples, derived from cell-type deconvolution analysis.

*DeconStatistics*

Left: statistical analysis comparing cell-type proportions for ASD v. controls in each region and each cell-type. Right: ANOVA with Tukey HSD p-values comparing cell-type proportions across all regions, stratified by ASD and Control. Only Bonferroni corrected significant p-values are shown.
